# Interspecies interactions alter the antibiotic sensitivity of *Pseudomonas aeruginosa*

**DOI:** 10.1101/2024.06.27.601049

**Authors:** C.I.M. Koumans, S.T. Tandar, A. Liakopoulos, J.G.C. van Hasselt

**Affiliations:** Leiden Academic Center for Drug Research, Leiden University, Leiden, The Netherlands; Microbiology, Department of Biology, Utrecht University, Utrecht, The Netherlands

## Abstract

Polymicrobial infections are infections that are caused by multiple pathogens, and are common in patients with cystic fibrosis (CF). Although polymicrobial infections are associated with poor treatment responses in CF, the effects of the ecological interactions between co-infecting pathogens on antibiotic sensitivity and treatment outcome are poorly characterized. To this end, we systematically quantified the impact of these effects on the antibiotic sensitivity of *Pseudomonas aeruginosa* for nine antibiotics in the presence of thirteen secondary cystic fibrosis-associated bacterial and fungal pathogens through time-kill assays. We fitted pharmacodynamic models to these kill curves for each antibiotic-species combination and found that interspecies interactions changing the antibiotic sensitivity of *P. aeruginosa* are abundant. Interactions that lower antibiotic sensitivity are more common than those that increase it, with generally more substantial reductions than increases in sensitivity. For a selection of co-infecting species, we performed pharmacokinetic-pharmacodynamic modelling of *P. aeruginosa* treatment. We predicted that interspecies interactions can either improve or reduce treatment response to the extent that treatment is rendered ineffective from a previously effective antibiotic dosing schedule and vice versa. In summary, we show that quantifying the ecological interaction effects as pharmacodynamic parameters is necessary to determine the abundance and the extent to which these interactions affect antibiotic sensitivity in polymicrobial infections.

**Importance:** In cystic fibrosis (CF) patients, chronic respiratory tract infections are often polymicrobial, involving multiple pathogens simultaneously. Polymicrobial infections are difficult to treat as they often respond unexpectedly to antibiotic treatment, which might possibly be explained because co-infecting pathogens can influence each other’s antibiotic sensitivity, but it is unknown to what extent such effects occur. To investigate this, we systematically quantified the impact of co-infecting species on antibiotic sensitivity, focusing on *P. aeruginosa*, a common CF pathogen. We studied for a large set co-infecting species and antibiotics whether changes in antibiotic response occur. Based on these experiments, we used mathematical modeling to simulate *P. aeruginosa*’s response to colistin and tobramycin treatment in the presence of multiple pathogens. This study offers comprehensive data on altered antibiotic sensitivity of P. aeruginosa in polymicrobial infections, serves as a foundation for optimizing treatment of such infections, and consolidates the importance of considering co-infecting pathogens.

## Introduction

Patients with cystic fibrosis (CF) suffer from chronic lung infections (1). Polymicrobial infections (PMIs), i.e., infections involving multiple microbial species simultaneously, are common in patients with CF (**Figure 1A**) (2–5). *Pseudomonas aeruginosa* is the most common pathogen in CF-PMIs in adults, but a variety of other microbial species has been found to co-infect (6, 7), potentially complicating CF treatment. For some of these co-infecting species, such as *Staphylococcus aureus, Haemophilus influenzae* and *Burkholderia cepacia*, the impact on the course of the infection has been well-established (8, 9). For other co-infecting species, such as *Achromobacter xylosoxidans, Streptococcus pneumoniae* and *Ralstonia mannitolilytica*, it remains unclear to what extent these organisms contribute to disease progression in chronic CF-associated respiratory tract infections (10–15).

**Figure 1.**
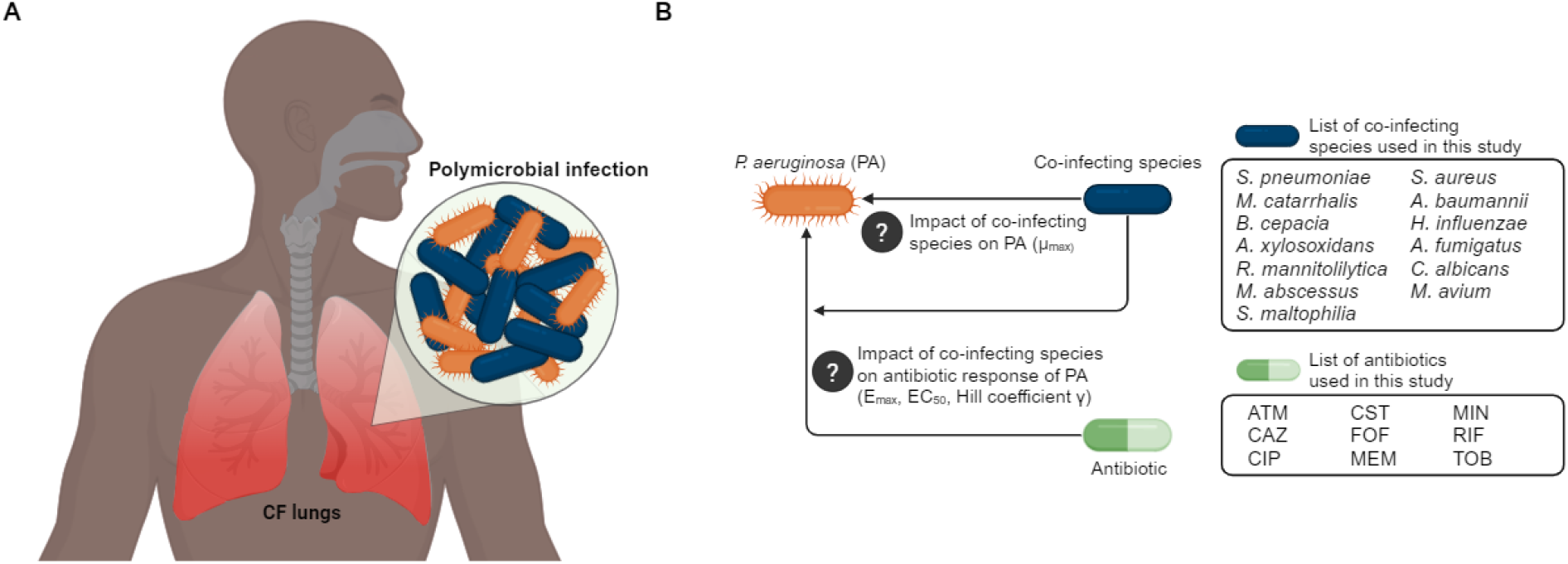
**A**. Polymicrobial infections in cystic fibrosis patients are common. **B**. A schematic representation of the routes of so far unexplored impact of co-infecting species in cystic fibrosis-polymicrobial infections and an overview of species and antibiotics used in this study. *ATM = aztreonam, CAZ = ceftazidime, CIP = ciprofloxacin, CST = colistin, FOF = fosfomycin, MEM = meropenem, MIN= minocycline, RIF = rifampicin, TOB = tobramycin, PA = P. aeruginosa*.

Antibiotic treatment of CF-PMIs is notoriously difficult. CF-PMIs are rarely fully cleared, requiring long-term antibiotic treatment to suppress exacerbations (16–18). PMIs can respond unexpectedly to antibiotic treatments. Treatment response may differ from what is expected from initial antibiotic sensitivity tests of single species of the CF-PMIs (19, 20). It is unclear if the presence of multiple pathogenic species in CF-PMIs alter antibiotic efficacy as compared to monomicrobial infections, and to what extent such effects should be considered in treatment guidelines. So far, limited *in vitro* data has shown that species in PMIs can interact with potential effects on antibiotic sensitivity (21–25).

Due to a lack of systematic data on these interspecies interactions, the general impact of such interactions on antibiotic treatment outcome in CF-PIMIs remains unknown. In this context, obtaining specific understanding of effects of interspecies interactions on antibiotic pharmacodynamics (PD) is essential. Determining changes in minimum inhibitory concentrations (MIC) is not sufficient to evaluate the potential impact of interspecies interactions on treatment response, as the MIC is a composite metric which combines changes in antibiotic sensitivity and growth rate, at one time point (26). In contrast, when expressing the impact of interspecies interactions as changes in pharmacodynamic (PD) parameters, the effect of interspecies interactions may be specifically attributed to specific PD parameters. These parameters include antibiotic sensitivity (EC50), maximum antibiotic effect (Emax), the sensitivity of pathogen kill rate (Hill slope), and changes in pathogen fitness in the absence of antibiotic (*μ*_*max*_) (27), which can then be used as part of pharmacokinetic-pharmacodynamic (PK-PD) analyses.

In this study we aimed to we systematically determine the impact of a large set of relevant CF-associated pathogens on the fitness and pharmacodynamics of *P. aeruginosa* for a range of antibiotics (**Figure 1B**). To this end, we cultured *P. aeruginosa* in medium conditioned by each secondary species separately as a proxy for the presence of a co-infecting species (**Figure 2A**). In this conditioned medium, we performed antibiotic time-kill studies for *P. aeruginosa* for different combinations of antibiotics and secondary species, which enabled assessment of changes in the PD response (**Figure 2B**). To evaluate the impact of interspecies interactions on antibiotic treatment schedules of *P. aeruginosa*, we performed pharmacokinetic-pharmacodynamic (PK-PD) modelling for selected antibiotic-secondary species combinations. Together, these results give guidance on the potential impact of interspecies interactions for antibiotic treatment strategies for CF-PMIs.

**Figure 2.**
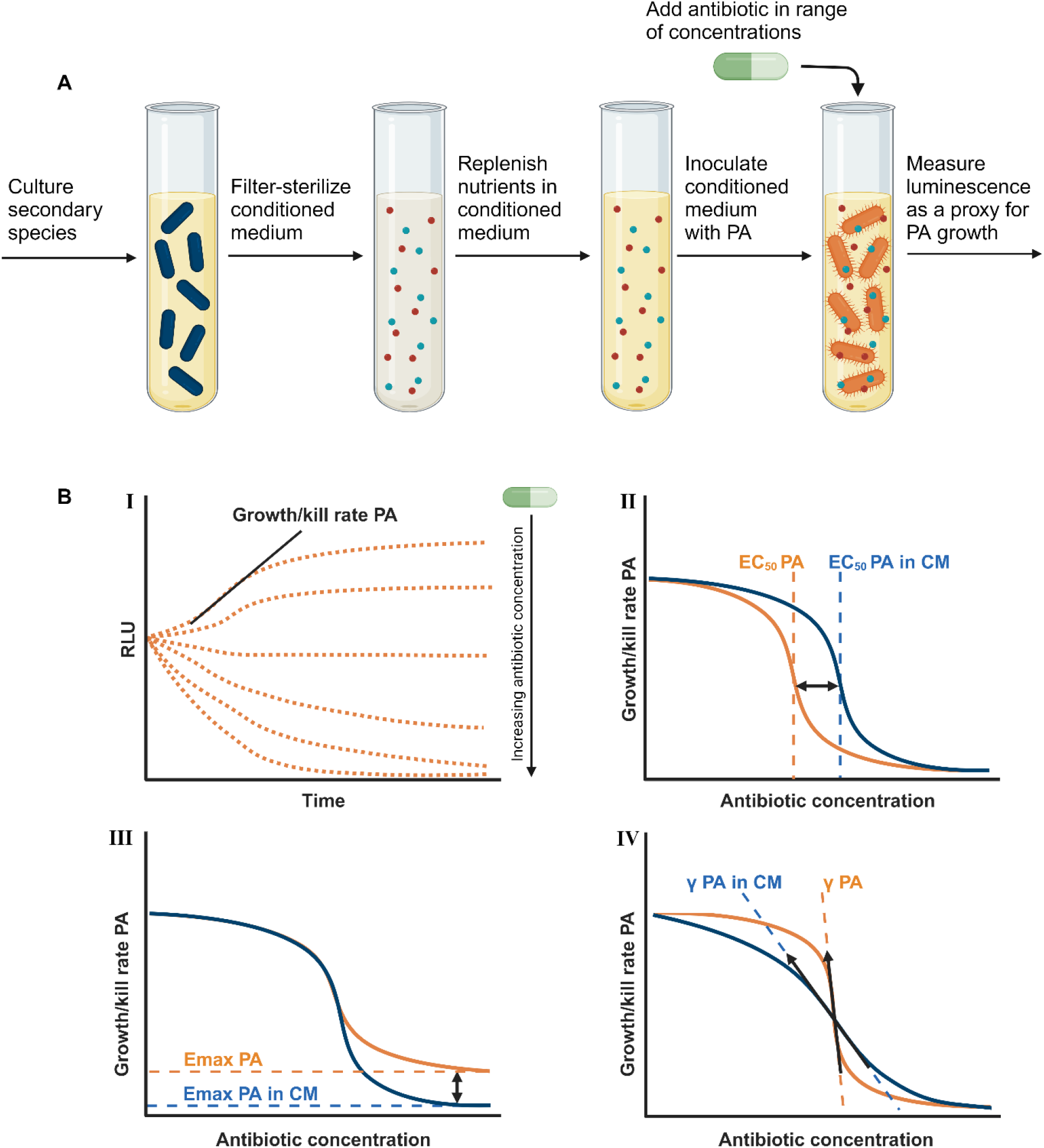
Schematic overview of experimental set-up and PD analysis. **A**. Preparation of conditioned medium and exposure of PA in conditioned medium to antibiotics. The secondary species was cultured in cation adjusted Mueller-Hinton broth allowing for the release of toxins and/or other metabolites inside the medium. Then the conditioned medium was filter-sterilized in order to remove the secondary species but not their toxins and/or other metabolites, and then nutrients were replenished. This replenished conditioned medium was used to culture *P. aeruginosa*, allowing us to determine the growth dynamics by measuring luminescence over time. **B**. I. Determination of μ_max_ by log-linear regression on exponential growth phase. II-IV. Visual representation of representative changes in PD parameters *EC*_50_, *E*_*max*_ and *γ* expressed as changes in Hill function, comparing conditioned and non-conditioned medium.

## Materials & Methods

### Strains

The focal pathogen strain used in this study was PAO1-Xen41, a bioluminescent *P. aeruginosa* strain encoding a single chromosomal copy of the *luxCDABE* operon (PerkinElmer). Strains representative of the 13 CF-associated secondary species that were used in the pairwise interaction assays together with *P. aeruginosa* PAO1-Xen41 included *Achromobacter xylosoxidans* DSM2402, *Acinetobacter baumannii* DSM30007, *Aspergillus fumigatus* DSM819, *Burkholderia cepacia* DSM7288, *Candida albicans* DSM1386, *Haemophilus influenzae* DSM44196, *Mycobacterium abscessus* DSM44196, *Mycobacterium avium* ATCC700898, *Moraxella catarrhalis* DSM9143, *Ralstonia mannitolilytica* DSM17512, *Staphylococcus aureus* DSM346, *Stenotrophomonas maltophilia* DSM21257, and *Streptococcus pneumoniae* DSM14377 (Leibniz Institute DSMZ-German Collection of Microorganisms and Cell cultures).

### Preparation of conditioned medium

The focal pathogen *P. aeruginosa* was cultured in medium conditioned by each secondary species (**Figure 2A**). The conditioned medium was used to approximate the presence of a co-infecting species in order to assess the contact-independent interaction between *P. aeruginosa* and each of the secondary species. Preparation of the conditioned medium started by culturing each secondary species on Mueller-Hinton agar (MHA) with species-specific supplements and culture conditions (**SI 1: Strains & Culture Conditions**). All secondary species were incubated at 37°C for 24 hours, with the exception of *M. abscessus* and *S. pneumoniae*, which were allowed to grow for 48 hours, and *M. avium*, which was allowed to grow for 5 days as they have longer incubation times. After incubation on agar medium, suspensions of 0.5 MacFarland (approximately 1.5 × 10^8^ colony forming units (CFU)/mL) were made in 0.9% saline. This suspension was then diluted down to 3000 CFU/mL. Multiple tubes containing 40 mL of cation-adjusted Mueller-Hinton broth (CAMHB) with species-specific supplements were inoculated with 1 mL of the 3000 CFU/mL cell suspension (**SI 1: Strains & Culture Conditions)**. All species were cultured at 37°C with agitation at 150 rotations per minute (rpm) for 48 hours, with the exception of *M. abscessus* and *M. avium*, which were cultured for 96 hours. Each culture tube was subsequently centrifuged (4654 g; 15 minutes) and filter-sterilized (0.22 µm) to remove secondary species from the conditioned medium. Conditioned medium nutrients were replenished by adding sterile 10x concentration CAMHB containing the respective species-specific medium supplements (10% v/v).

### Antibiotics

Antibiotic preparation was done in advance of all the experiments according to the manufacturers’ recommendation with aztreonam, ceftazidime, ciprofloxacin, colistin, fosfomycin, meropenem, minocycline, rifampicin and tobramycin being dissolved in their respective solvents, aliquoted and stored at the specific storage conditions required (**SI 2: Antibiotics**). Before each experiment, antibiotic stock solutions were diluted in the conditioned medium required to obtain the desired range of concentrations (**SI 2: Antibiotics**).

### Time-kill experiments

Time-kill experiments of *P. aeruginosa* (**Figure 2A**) were performed for all antibiotics in medium conditioned with each secondary species. The time-kills were performed in 96-well plates (250 µL culture volume) with a *P. aeruginosa* starting cell density of approximately 5 × 10^6^ CFU/mL. Time-kill plates were incubated aerobically at 37°C with agitation at 150 rpm. The number of viable bacterial cells over time was estimated by measuring luminescence signal and relative light units (RLU) hourly for 20 hours (BMG Fluostar Omega; gain 3800; interval 1.36 seconds).

### Pharmacodynamic analysis

We first computed the maximum growth rate for each specific combination of an antibiotic, secondary species, and drug concentration by log-linear regression on the exponential growth/kill phase. For this purpose, we developed a phase selection script, which automatically determines the exponential growth/kill phase (**SI 3: Visualization Phase Selection Script**).

In order to quantify the PD parameters that display the antibiotic sensitivity of *P. aeruginosa*, concentration-effect relationships (Equation 1) were fitted for the individually estimated growth rates against the corresponding antibiotic concentrations for each co-infecting species-antibiotic combination:

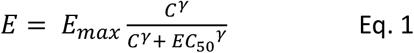

Here, *E* represents the growth/kill rate of *P. aeruginosa, C* represents the drug concentration, *E*_*max*_ the maximum effect, *EC*_50_ the drug concentration to achieve a half-maximum drug effect and the Hill coefficient *γ* represents the steepness of the dose-response curve. The fitting procedure was performed using the R package ‘drc’ (28). The effect of secondary pathogen on PD parameters *E*_*max*_, *EC*_50_ and *γ* (**Figure 2B**) were quantified as its fold-change (FC) to its respective value estimated in non-conditioned medium. We considered log2(*FC*) > 0.5 to indicate an increase in the specific PD parameter and log2(*FC*) > −0.5 to indicate a decrease. The impact of co-infecting pathogens on the fitness of *P. aeruginosa* was determined by dividing the maximum growth rate *μ*_*max*_ in the absence of drugs in conditioned medium by the *μ*_*max*_ in the absence of drugs in a non-conditioned medium, obtaining the FC and determining whether there was an increase or decrease in fitness in the same way as the PD parameters. All *μ*_*max*_ and PD parameter values can be found in **SI 4: Mumax Values** and **SI 5: PD Values**.

### Mathematical pharmacokinetic-pharmacodynamic modelling

Mathematical PK-PD model-based analysis was performed to evaluate the impact of selected secondary species on the clinical outcome of *P. aeruginosa* treatment with either colistin or tobramycin. First, published population PK models for colistin and tobramycin were implemented to describe drug concentration in the lung over the course of antibiotic treatment, using the standard treatment regime from these studies (29, 30). Simulated colistin dosing involved a 160 mg loading dose (inhalation) followed by 160 mg maintenance doses (intravenous; 30 minutes infusion) with an interval of 8 hours. Simulated tobramycin dosing involved a constant intravenous (bolus) administration of 139.65 mg tobramycin with an 8-hours interval. We simulated the treatment for a typical individuals with the pharmacokinetic clearance parameter were increased with one standard deviation of the mean from the patient population for colistin, reflecting the sub-population of patients with increased drug clearance and therefore reduced colistin exposure, for tobramycin the standard clearance parameters were used. A tobramycin lung-plasma partition coefficient of 0.12 was used to calculate lung concentration based on the plasma concentration (Eq. 2)(31). This conversion was not necessary for colistin, as the model already provided the lung PK directly.

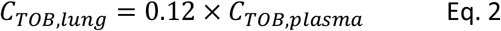

A population-limited growth model was used to describe the growth behavior of *P. aeruginosa* in different antibiotic conditions (Eq. 3).

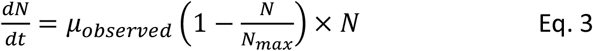

Here, *N*_*max*_ is the maximum cell density of *P. aeruginosa* in the epithelial lining fluid (ELF) of the lung and was fixed to 9 × 10^10^ CFU/mL based on the observed carrying capacity of our the experimental system. *N* is the number of CFU/mL of *P. aeruginosa* at a given time in the simulation and is set at 5 × 10^6^ CFU/mL at time 0 as this was the starting cell density of our time-kill experiments. The observed growth rate *μ*_*observed*_ was calculated using Eq. 4.

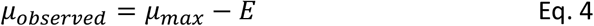

Bacterial growth kinetic- and drug effect parameters estimated from the experimental data were used to determine the effect of the selected co-infecting species on the drug response of *P. aeruginosa* expressed as changes in *μ*_*observed*_ . To evaluate the impact of co-infecting species, we selected a number of species with a variable effect on PD parameters to illustrate their potential impact. All PK-PD parameters, details of antibiotic dosing schedules, and other variable values can be found in **SI 6: PK-PD Simulation Parameter Values**.

## Results

### Co-infecting species affected the *μ*_*max*_ of *P. aeruginosa* in absence of antibiotics

We plotted the measured RLU over time to obtain growth/kill curves of P. aeruginosa in contact-independent interaction with each of the co-infecting species. We did this in absence of antibiotics and in presence of multiple concentrations of each antibiotic. All raw growth/kill curves that we obtained this way can be found in **SI 7: Raw Growth Curves**. We first studied the effect of species interactions under absence of antibiotic treatment on bacterial growth rates, which may be important contributors to microbial composition of PMIs as well as the response to antibiotic treatment. To determine what the impact on CF-associated secondary species is on the *μ*_*max*_ of *P. aeruginosa*, we compared the *μ*_*max*_ of *P. aeruginosa* in non-conditioned medium to the *μ*_*max*_ in conditioned medium, in the absence of any antibiotic. We showed that co-infecting secondary species were able to significantly alter the *μ*_*max*_ of *P. aeruginosa* (**Figure 4D**). Specifically, *A. xylosoxidans, A. baumannii, B. cepacia, M. abscessus* and *S. aureus* caused a decrease, whereas *A. fumigatus* and *H. influenzae* caused an increase in *μ*_*max*_. A decrease in *μ*_*max*_ indicates that growth rate of *P. aeruginosa* is lower when the secondary species is present, whereas an increase indicates that the growth rate is higher, meaning that the population size of *P. aeruginosa* in a PMI might differ depending on which secondary species are present.

### Co-infection by secondary species altered the pharmacodynamics of antibiotics against *P. aeruginosa*

To assess the impact of a secondary species on antibiotic pharmacodynamic parameters for *P. aeruginosa*, we determined the growth/kill rate of *P. aeruginosa* for a range of antibiotics and antibiotic concentrations in medium conditioned by each of the CF-associated species. We then derived concentration-effect curves by plotting the growth/kill rates over the corresponding antibiotic concentrations. This allowed us to compare concentration-effect curves for conditioned and non-conditioned medium for the same antibiotic-secondary species combination. All co-infecting species are capable of altering each of the pharmacodynamic parameters, but it depends on the antibiotic which PD parameters are affected and whether they are increased or decreased. To illustrate how the changes in growth/kill curves correspond to changes in the concentration-effect curves, we selected representative examples for meropenem (**Figure 3A-B**) and ciprofloxacin (**Figure 3C-D**). We observed similar changes for a variety of antibiotic-secondary species combinations (**SI 8: Concentration Effect Curves**). For instance, the growth rate of *P. aeruginosa* in monomicrobial infection was decreased compared to co-infection with either *A. baumannii, A. xylosoxidans*, or *S. maltophilia* during treatment with 2 mg/L meropenem (**Figure 3A**), visible in the corresponding concentration-effect curves (**Figure 3B**) as a shift to the right, denoting an increase in *EC*_50_ . Conversely, the growth rate of *P. aeruginosa* in monomicrobial infection was increased compared to co-infection with either *A. xylosoxidans, B. cepacia* or *S. aureus* during treatment with 0.063 mg/L ciprofloxacin (**Figure 3C**), visible as a shift to the left (**Figure 3D)**, denoting a decrease in *EC*_50_ . While for meropenem this does not appear to result in a large change in the steepness of the concentration-effect curves (Hill coefficient *γ*), for ciprofloxacin, the presence of a co-infecting species leads to a reduction in steepness compared to the control. Changes in *E*_*max*_ are also visible as an upward or downward shift in the lower asymptote for both antibiotics. These examples illustrate how the PD parameters used cover the full range of the antibiotic response.

**Figure 3.**
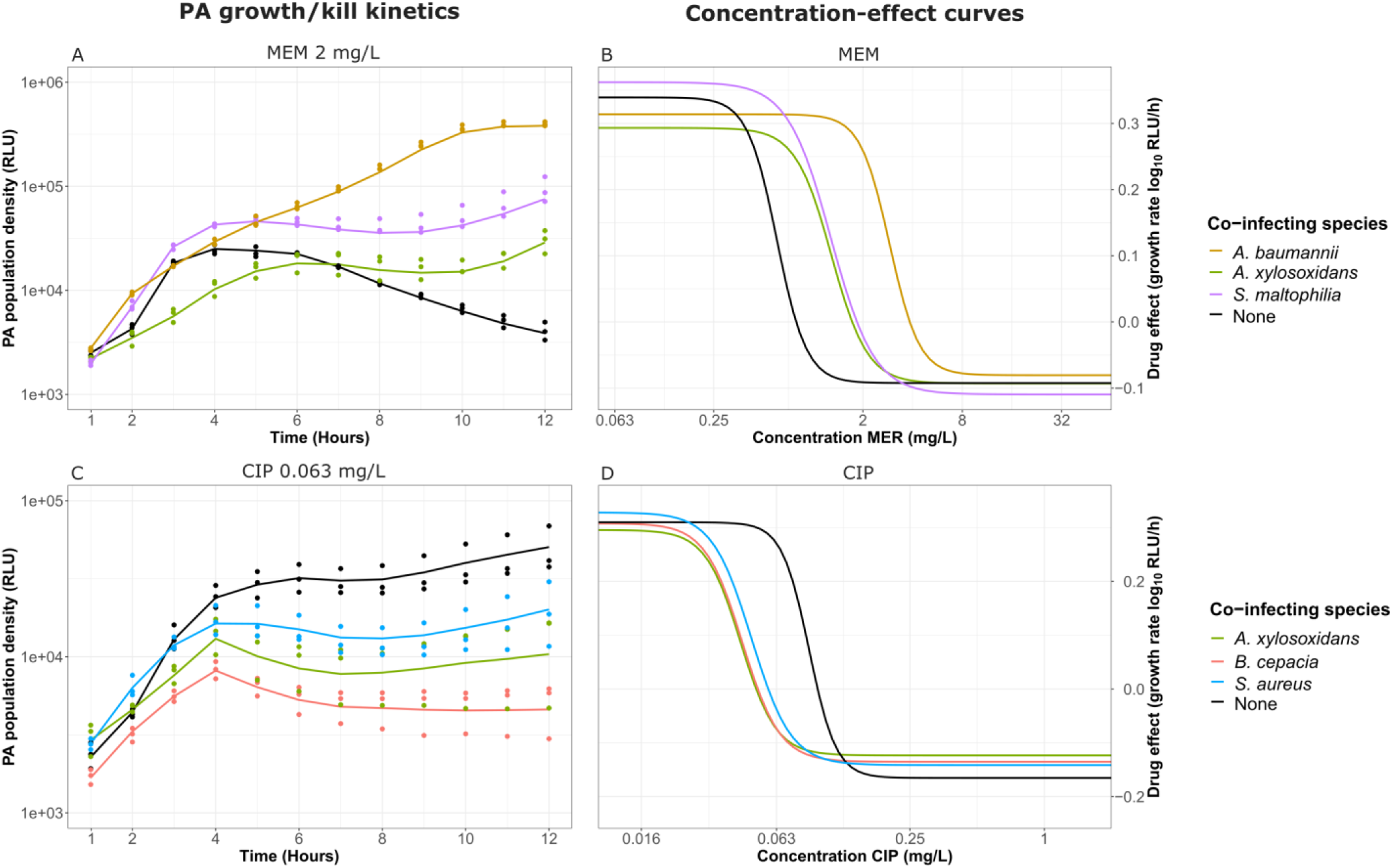
Representative examples of the translation of growth/kill kinetics to the concentration-effect curves. **A**. *P. aeruginosa* growth/kill kinetics in the presence of 2 mg/L meropenem in monomicrobial infection and co-infection. **B**. Concentration-effect curves of *P. aeruginosa* in the presence of meropenem in monomicrobial infection and co-infection. **C**. *P. aeruginosa* growth/kill kinetics in the presence of 0.063 mg/L ciprofloxacin in monomicrobial infection and co-infection. **D**. Concentration-effect curves of *P. aeruginosa* in the presence of ciprofloxacin in monomicrobial infection and co-infection. MEM = meropenem, CIP = ciprofloxacin.

### The presence of a co-infecting species leads more often to a reduction in antibiotic sensitivity of *P. aeruginosa*

To determine the impact of the interspecies interactions during CF on the antibiotic response of *P. aeruginosa*, we first focused on the *EC*_50_. When a secondary species increases the *EC*_50_ of an antibiotic, it means that a higher concentration of this antibiotic is needed to reach the same impact on *P. aeruginosa* compared to *P. aeruginosa* in monomicrobial infection. This, in turn, indicates that the species interaction leads to a reduction in antibiotic sensitivity for *P. aeruginosa*. On the contrary, when a secondary species decreases the *EC*_50_, this corresponds to an increase in antibiotic sensitivity for *P. aeruginosa*. Our results demonstrate that all secondary co-infecting species lead to a change in *EC*_50_ for the majority of the antibiotics tested, except for aztreonam, where none of the secondary species changed the *EC*_50_ (**Figure 4A**). For colistin, interactions only lead to an increase in *EC*_50_, whereas for the other antibiotics, the impact of secondary species was bidirectional. Depending on the antibiotic, all secondary species caused either increases or decreases in *EC*_50_, except for *A. baumannii, H. influenzae, M. avium* and *M. catarrhalis*, which only found to increase the *EC*_50_. The impact of *R. mannitolilytica* and *S. aureus* was remarkably similar as both increased the *EC*_50_ for minocycline and fosfomycin and decreased the *EC*_50_ for ciprofloxacin and tobramycin. *S. maltophilia* and *A. xylosoxidans* most often caused a change in *EC*_50_, for 7 out of the 9 antibiotics tested. The direction of the impact of *S. maltophilia* and *A. xylosoxidans* on the sensitivity of these antibiotics was also similar, with the exception of tobramycin where *S. maltophilia* did not alter the *EC*_50_ and ceftazidime where *A. xylosoxidans* did not alter the *EC*_50_. Overall, we showed that interactions that lead to an increase in *EC*_50_ of *P. aeruginosa* were more commonly observed (41.9%) than interactions that lead to a decrease in *EC*_50_ (12.8%) indicating that the antibiotic sensitivity of *P. aeruginosa* was more often reduced than increased in co-infections with other secondary CF-associated species (**SI 9: Distribution PD parameters**).

**Figure 4.**
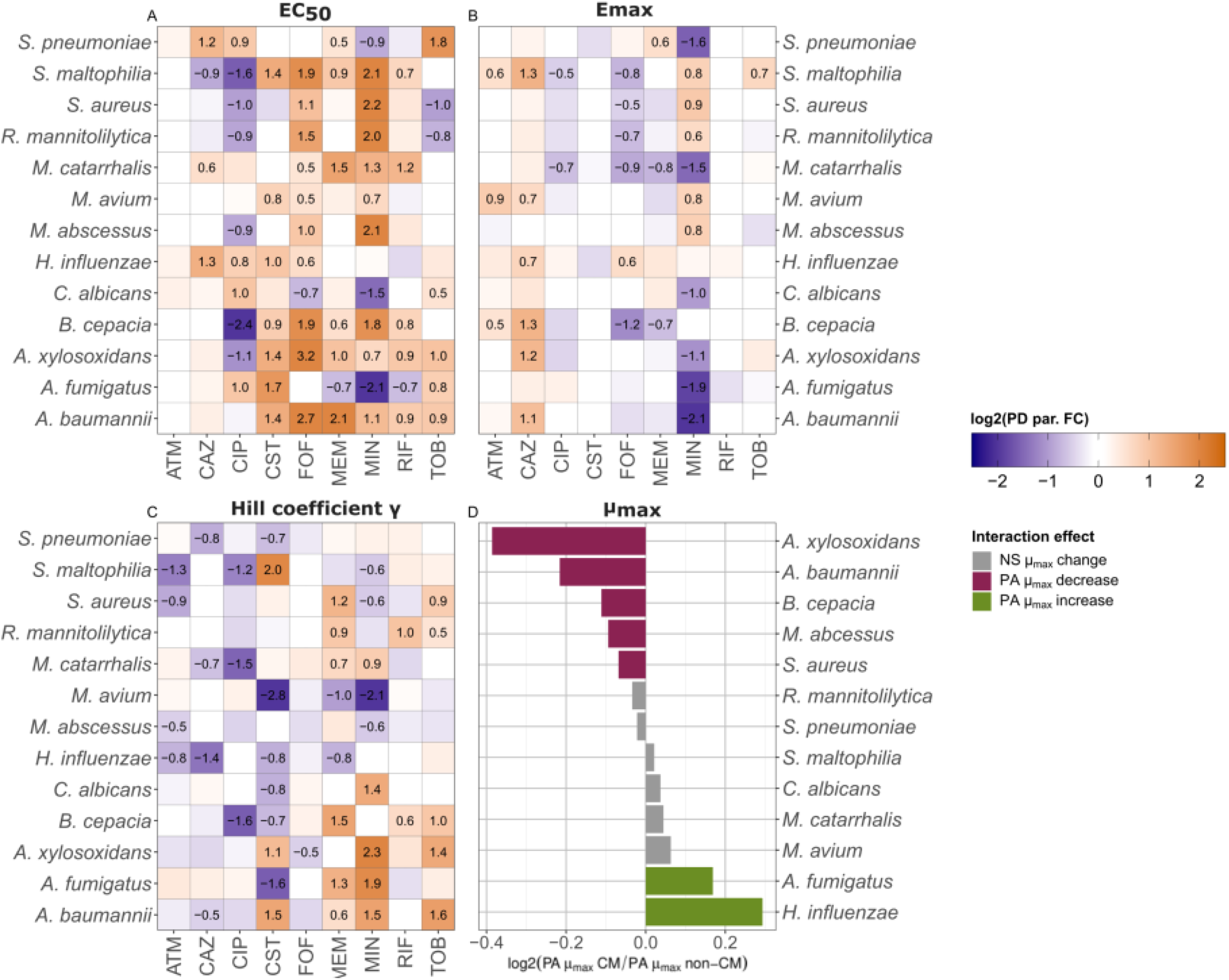
Changes in PD parameters *EC*_*50*_, *E*_*max*_ and Hill coefficient *γ* and *μ*_*max*_ in the presence of a CF-associated co-infecting species. **A**. Change in PA *EC*_50_. **B**. Change in PA *E*_*max*_ . **C**. Change in PA Hill coefficient *γ*. Cells in heatmaps A-C are colored when log2(*FC*) > 0.15 or log2(*FC*) < −0.15. Cells in heatmaps A-C display the value of the parameter changes when log2(*FC*) > 0.5 or log2(*FC*) < −0.5. **D**. Change in PA *μ*_*max*_ in the absence of antibiotics. Significance (p<0.01) was determined by performing a two-samples unpaired t-test for comparing the *μ*_*max*_ of *P. aeruginosa* in conditioned medium (CM) and non-conditioned medium (non-CM). NS = non-significant. Purple means significant decrease and green means significant increase in PA *μ*_*max*_ . ATM = aztreonam, CAZ = ceftazidime, CIP = ciprofloxacin, CST = colistin, FOF = fosfomycin, MEM = meropenem, MIN= minocycline, RIF = rifampicin, TOB = tobramycin.

### Direction of interaction-mediated changes in *E*_*max*_ and Hill coefficient *γ* were distributed more evenly than changes in *EC*_50_

In addition to changes in the *EC*_50_ of *P. aeruginosa* due to its interaction with CF-associated co-infecting species, we observed changes in the *E*_*max*_ and Hill coefficient *γ* highlighting the impact of these co-infecting species on the antibiotic response of *P. aeruginosa*. We showed that these PD parameters changed independently from each other and from the *EC*_50_. Among others, we specifically showed that the *EC*_50_ for rifampicin and colistin were altered by multiple interspecies interactions, but there were no changes in *Emax* for these drugs. The opposite was the case for aztreonam, where there were no changes in the *EC*_50_, but there were interaction-mediated changes in *E*_*max*_ and Hill coefficient *γ*. The distribution of the directions of the interspecies effects was also different for the *E*_*max*_ and Hill coefficient *γ* compared to the *EC*_50_. Interactions that increased the *E*_*max*_ (14.5%) show that the maximum killing by the antibiotic increases during treatment, and vice versa for interactions that decreased the *E*_*max*_ (12.8%) (**Figure 4B**). *S. maltophilia* most often altered the *E*_*max*_, by either increasing and decreasing it. *M. catarrhalis* was the only species that exclusively decreased the *E*_*max*_. Interactions that increased the Hill coefficient *γ* (17.9%) show a stronger killing effect if antibiotic concentration increases during treatment. (**Figure 4C**). In contrast, interactions that decreased the Hill coefficient *γ* (20.5%) mean that a larger increase in antibiotic concentration is needed to increase the killing effect in co-infection compared to monomicrobial infection. Taken together, we showed that changes in PD parameters *E*_*max*_ and Hill coefficient *γ* were more evenly distributed between increases and decreases than the changes in *EC*_50_ (**SI 9: Distribution PD parameters**).

### Co-infecting species could affect the treatment outcome of *P. aeruginosa* with tobramycin and colistin

To determine whether the observed impact of the co-infecting species on the PD parameters and *μ*_*max*_ affects the treatment outcome of *P. aeruginosa* in a CF-PMI, we simulated the effect of these observed changes on the *P. aeruginosa* eradication during treatment of a polymicrobial CF-lung infection. We focused on tobramycin and colistin, which are clinically relevant antipseudomonal antibiotics with narrow therapeutic windows (**Figure 5**). The chosen array of co-infecting species caused both increasing and decreasing effects on the PD parameters and specifically on the *EC*_50_, allowing us to explore the potential effects of interspecies interactions during polymicrobial infections on antibiotic treatment. Colistin treatment (**Figure 5A)** in monomicrobial infection lead to eradication at the end of the simulation (**Figure 5B**) whereas tobramycin treatment (**Figure 5C)** in monomicrobial infection lead to a stable number of P. aeruginosa CFU/mL (**Figure 5D**). In co-infection with *A. baumannii* or *A. xylosoxidans, P. aeruginosa* treatment failed with either antibiotic as the size of the initial infecting population of *P. aeruginosa* increased over the course of the simulated lung infection despite the antibiotic treatment. In contrast, *P. aeruginosa* was eradicated in co-infection with *S. aureus* treated with either antibiotics. In the case of *P. aeruginosa* treated with colistin in co-infection with *S. aureus*, it is remarkable that even for a relatively small change in antibiotic sensitivity (log2(*FC EC*_50_) = −0.4) the outcome of treatment is altered compared to monomicrobial infection. For tobramycin, we also showed that co-infection with *R. mannitolilytica* is able to lead to eradication with *P. aeruginosa*. Collectively, these results show that the impact of interspecies interactions on the antibiotic response of *P. aeruginosa* could alter treatment outcomes to the extent that treatment is rendered ineffective.

**Figure 5.**
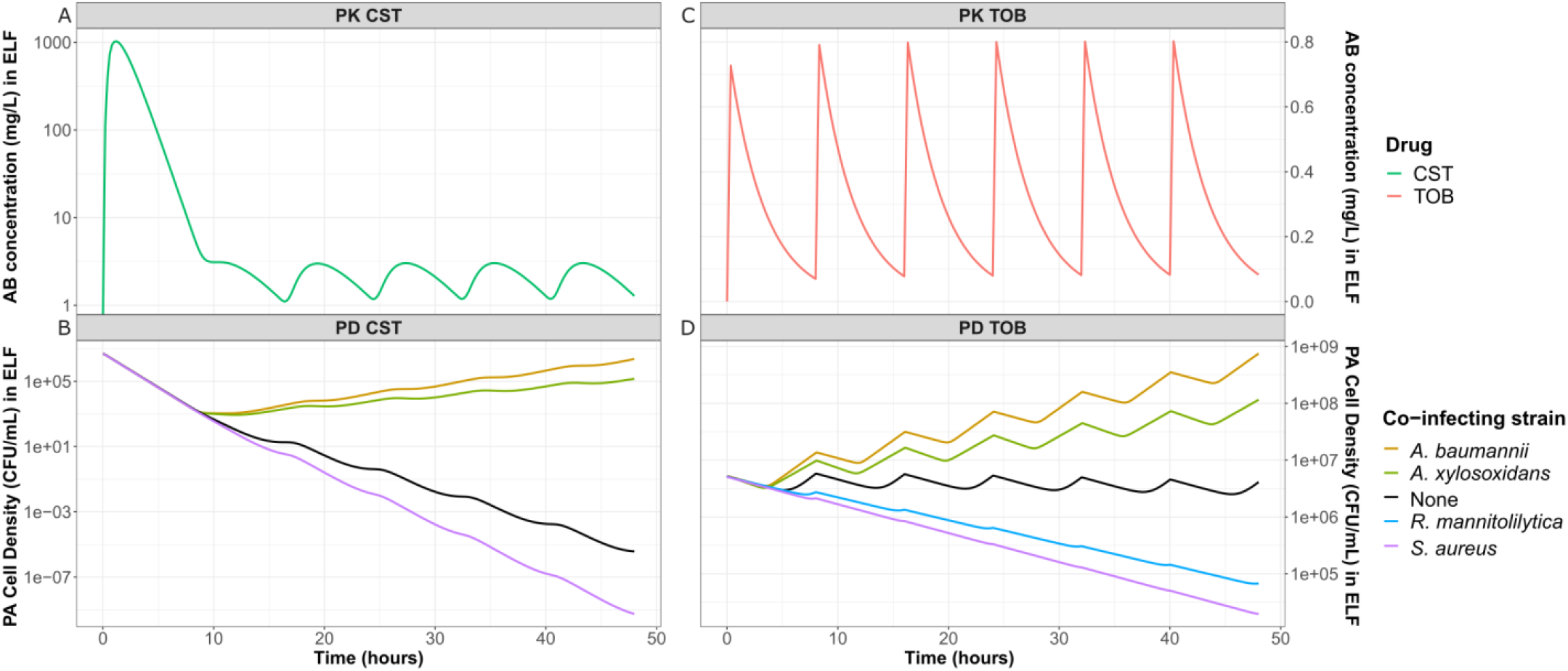
PK-PD simulation of colistin and tobramycin treatment of *P. aeruginosa* alone or in co-infection with a CF-pathogen. **A**. PK simulation of colistin (CST) concentration in epithelial lining fluid (ELF). **B**. PD simulation of *P. aeruginosa* exposed to colistin in ELF. C. PK simulation of tobramycin (TOB) concentration ELF. D. PD simulation of *P. aeruginosa* exposed to tobramycin in ELF. Starting cell density in the ELF of *P. aeruginosa* is 5 × 10^6^ CFU/mL for both tobramycin and colistin.

## Discussion

Chronic lung infections in CF patients are a characteristic example of polymicrobial infections (1), where contact-dependent and -independent pathogen-pathogen interactions are expected to alter the behaviors of the co-infecting species (32–34). In this study, we established an experimental framework and a subsequent analysis pipeline to specifically investigate the contact-independent impact of any CF-associated secondary species on critical parameters for the antibiotic eradication of *P. aeruginosa* while approximating a CF-PMI. In particular, we performed high-throughput time-kill assays of *P. aeruginosa* in medium previously conditioned separately by each of the CF-associated secondary species for an extensive range of antibiotics and antibiotic concentrations, and we systematically determined changes in antibiotic response (PD parameters) and population size (*μ*_*max*_) of *P. aeruginosa*.

Our data showed extensive impact of the secondary species on *P. aeruginosa* PD parameters and/or *μ*_*max*_. Given that we replenished the conditioned medium with nutrients after filtering out the secondary species, this observed contact-independent impact on *P. aeruginosa* is likely not due to nutrient depletion but due to metabolic by-products of these secondary species secreted in the conditioned medium. We showed that all secondary species included in our study are able to affect the antibiotic response of *P. aeruginosa* in an antibiotic-dependent manner and that a decrease was more common than an increase in sensitivity of *P. aeruginosa* (*i*.*e*., an increase in *EC*_50_). This was the case for the majority of the antibiotics to which we exposed *P. aeruginosa*, with the exception of aztreonam sensitivity where we observed no change, and ciprofloxacin sensitivity where an increase in sensitivity was more commonly observed. Similarly, interspecies interactions that occur in urinary tract infections (UTIs) were previously found to most commonly decrease the sensitivity of bacterial pathogens in UTIs against trimethoprim-sulfamethoxazole and nitrofurantoin (35). In addition to the changes in antibiotic sensitivity, we showed that in absence of antibiotics, secondary species could either increase or decrease the *μ*_*max*_ of *P. aeruginosa*. Overall, we indicated that a decrease in sensitivity was more commonly observed, which is in accordance with the notion that interspecies interactions are more often of a competitive than a cooperative nature (35, 36).

Direct comparison with previous findings is difficult, as the limited number of earlier studies employed a range of methods to approximate interspecies interactions. Studies differ from each other in their choice of isolates, use different readouts and expose the species to each other through several methods in different lifestyles, such as direct co-culture or conditioned medium in planktonic or biofilm culture (37–42). These differences in experimental set-up might be the reason for contradicting study results. For example, one study found that 10% medium conditioned by a clinical isolate of *S. aureus* decreased tobramycin sensitivity for the reference strain PAO1 and clinical *P. aeruginosa* strains isolated from children with CF for which previous antibiotic eradication therapy had failed (43). In contrast, a different study showed that planktonic and biofilm co-cultures of *S. aureus* ATCC25923 and *P. aeruginosa* in the presence of lung epithelial cells increased tobramycin sensitivity for three *P. aeruginosa* reference strains, but not for PAO1 (44). In our study, where we use a conditioned medium approach and a planktonic lifestyle, we found that *S. aureus* (DSM346) increases the tobramycin sensitivity of *P. aeruginosa* (PAO1-Xen41). These examples highlight the relevance of applying a high-throughput standardized workflow to enable the systematic exploration of the impact of interspecies interactions on *P. aeruginosa’s* antibiotic response.

Other studies determining the antibiotic response of *P. aeruginosa* in presence of other co-infecting species used different experimental approaches, such as a direct co-culture, or determine the response of *P. aeruginosa* in biofilm models (23, 40, 45, 46). For many of these experimental approaches, it can often be challenging to obtain time course data in a high throughput fashion, meaning the output of these studies is often for one antibiotic concentration at a single time point, or time course data can only be obtained in a low throughput fashion. As our goal was to determine in detail the antibiotic concentration-effect profiles of *P. aeruginosa* in order to perform PD analysis, we decided to focus on measuring bacterial growth over time and for many antibiotics and antibiotic concentrations. This makes a biofilm or direct co-culture experimental model unsuitable for our study aims, where our conditioned medium approach does meet the study requirements.

Our specific focus on identifying effects of species interactions on PD parameters and not MIC is important, as this allows to obtain an understanding at the pharmacological mode of action of interspecies interactions. In addition, depending on the specific PD parameter which is altered, different adjustments of the antibiotic dosing schedules need to be implemented in order to improve treatment, underlining, in turn, the need for pharmacodynamic analysis based on antibiotic time kill-curves instead of lump sum methods such as MIC determination (26, 27). In our study, all species were found to alter the antibiotic response of *P. aeruginosa* for one or more of the PD parameters tested in an antibiotic-dependent manner. For some of these species, such as *A. baumannii, A. xylosoxidans* and *S. maltophilia*, the impact on the severity of lung infections in CF is poorly understood (47, 48). Our results suggest that those species can alter the antibiotic response of *P. aeruginosa* during treatment, thereby contributing to the understanding of how CF-PMIs progress over time.

To further evaluate the potential clinical implications of interspecies interactions on antibiotic treatment regimens, we simulated antibiotic treatment of *P. aeruginosa* with colistin or tobramycin under presence of different secondary species, including *A. baumannii, A. xylosoxidans, S. aureus* or *S. maltophilia*. Our PK-PD simulations showed that the identified alterations in PD parameters induced by species interactions could have a clinically relevant impact, either potentiating treatment or leading to treatment failure. This is clinically relevant as colistin and tobramycin have narrow therapeutic windows, making the risk of underdosing likely when a pathogen-pathogen interaction results in decreased antibiotic sensitivity for *P. aeruginosa* (49, 50). Underdosing, in turn, not only leads to treatment failure, but may also lead to the development of antibiotic resistance (51). On the other hand, when a pathogen-pathogen interaction results in increased *P. aeruginosa* sensitivity, this could mean that less antibiotic is necessary to obtain treatment success which could be beneficial for patients by minimizing the toxic effect of colistin and tobramycin. The use of PKPD modeling strategies such as demonstrated for these case studies offer an important tool to help further translate in vitro pharmacodynamic data to the clinical situation and ultimately also offers the flexibility to further incorporate specific pharmacodynamic mechanisms (52, 53).

In conclusion, our study provide a comprehensive quantitative overview on interspecies interaction effects on the pharmacodynamic response of *P. aeruginosa* in the presence of CF-associated co-infecting species, and the potential to further translate such interaction effects to clinical dosing schedules through the use of PK-PD modeling. Our analyses demonstrates that identified interaction effects have the potential to alter antibiotic treatment outcomes, consolidating the relevance of interspecies interactions on the antibiotic treatment of CF-patients with PMIs. Overall, our study provides the foundation for further studies on the role of interspecies interactions to optimize antibiotic treatment of CF-PMIs.

## Acknowledgements

This research received no specific funding from any agency in the public, commercial, or not-for-profit sectors. We would like to thank Maik Kok for his contributions to fruitful discussions about the data analysis and the phase selection script.

## Data availability paragraph

The raw data of the time-kill curves is available upon reasonable request through the corresponding author.

