## Supplementary figures and images for "Interspecies interactions alter the antibiotic sensitivity of *Pseudomonas aeruginosa*"

### SI_3_Visualization_Phase_Selection_Script.pdf

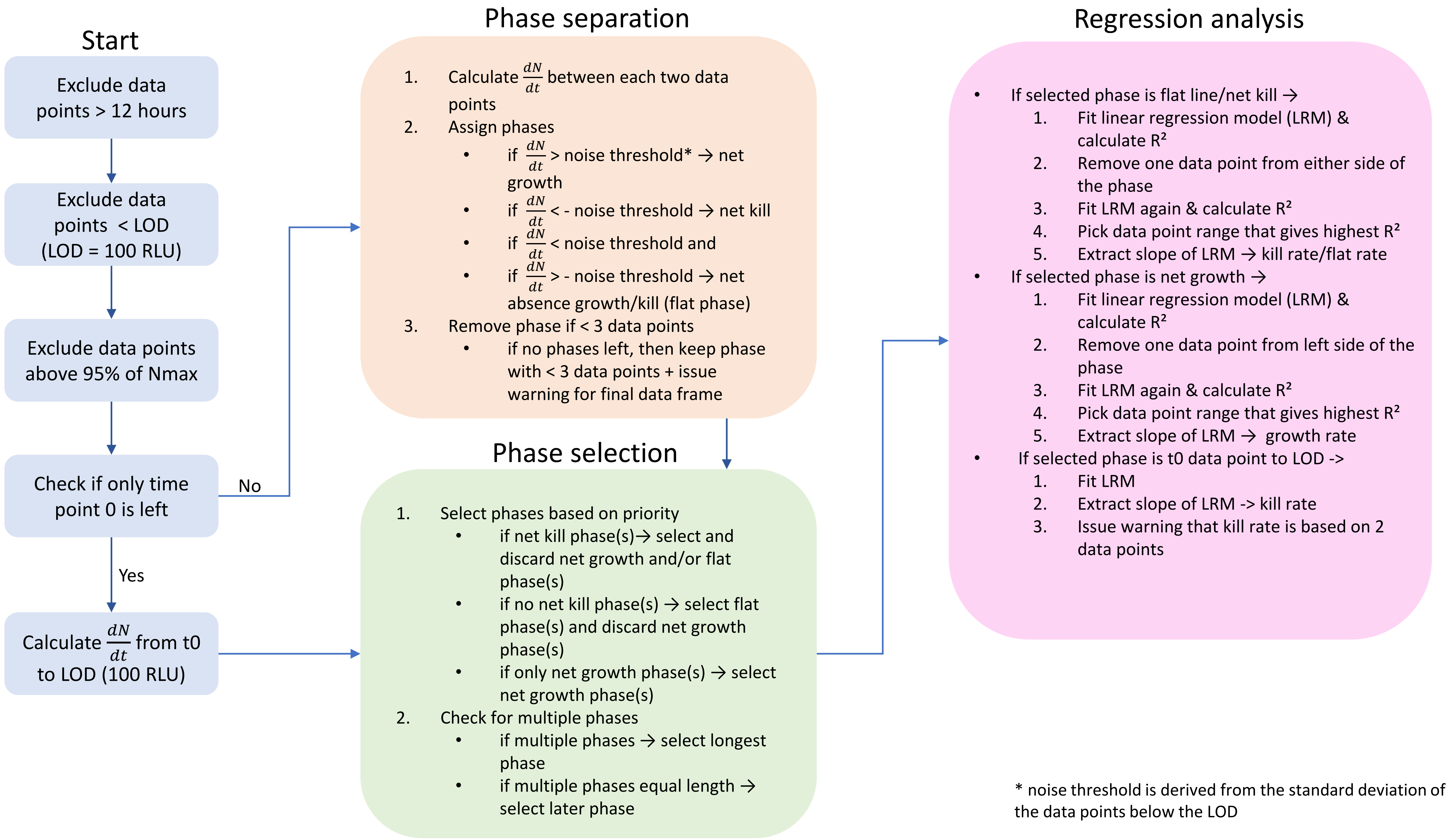

### SI_7_Raw_Growth_Curves.pdf

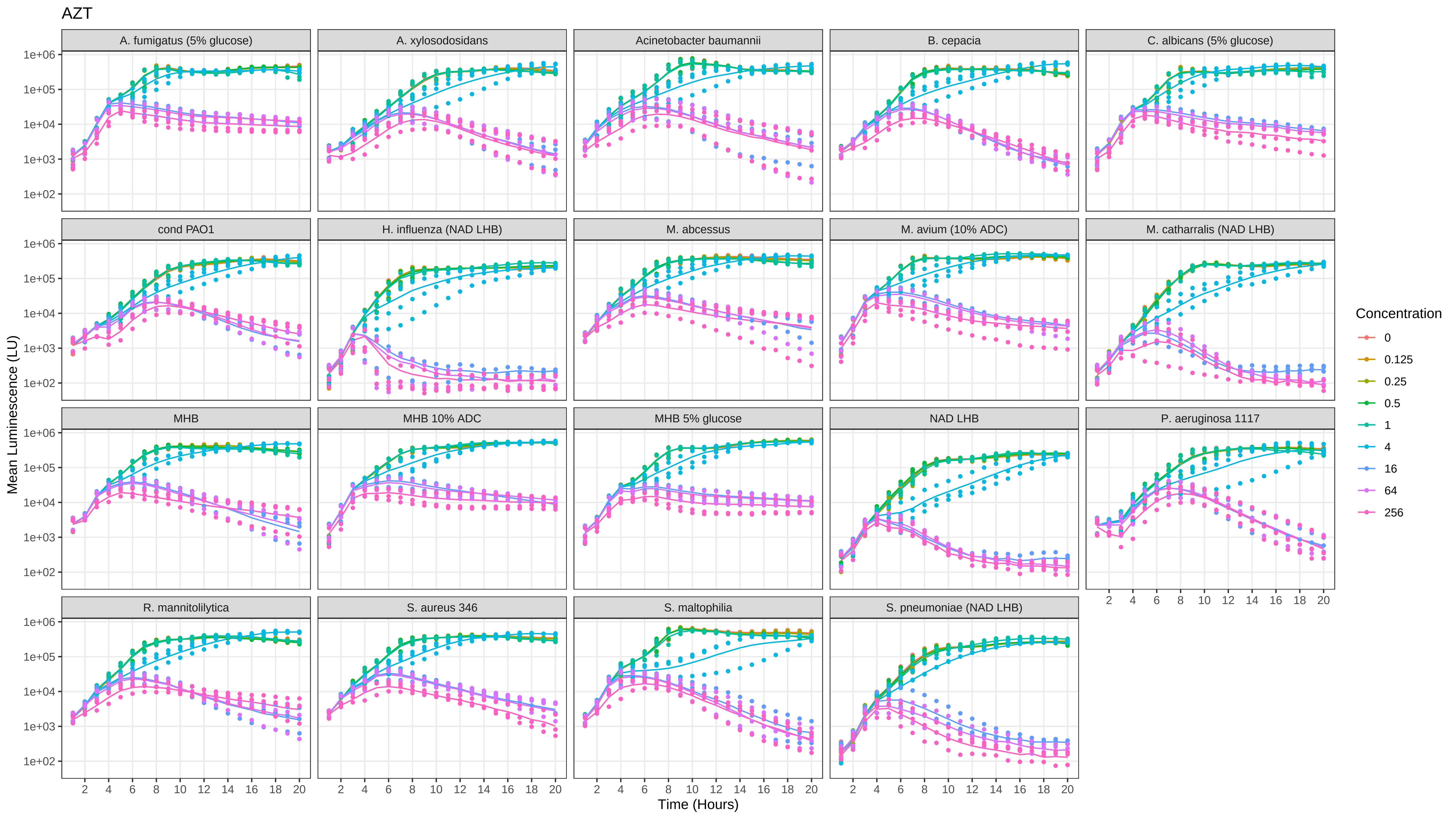

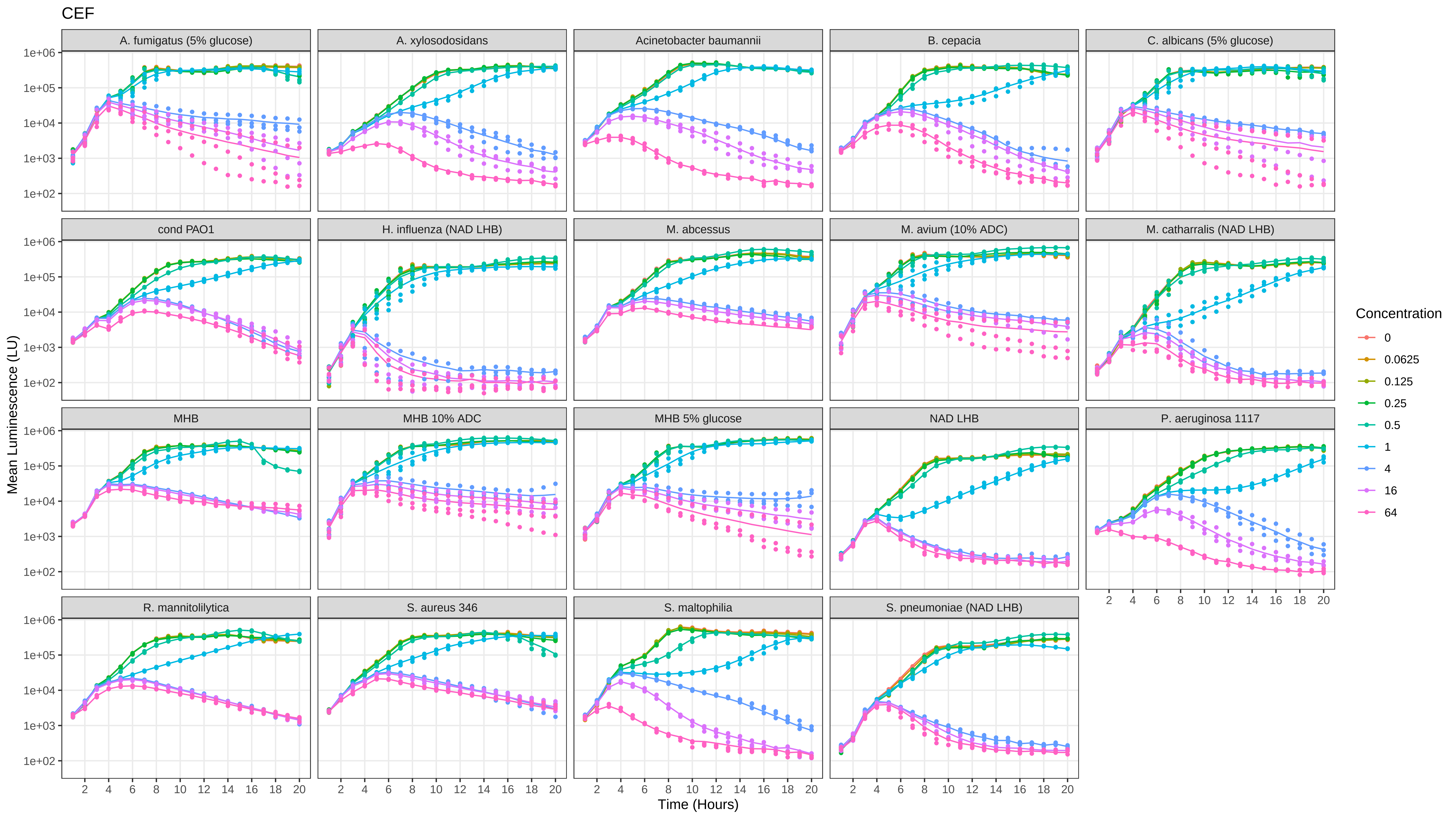

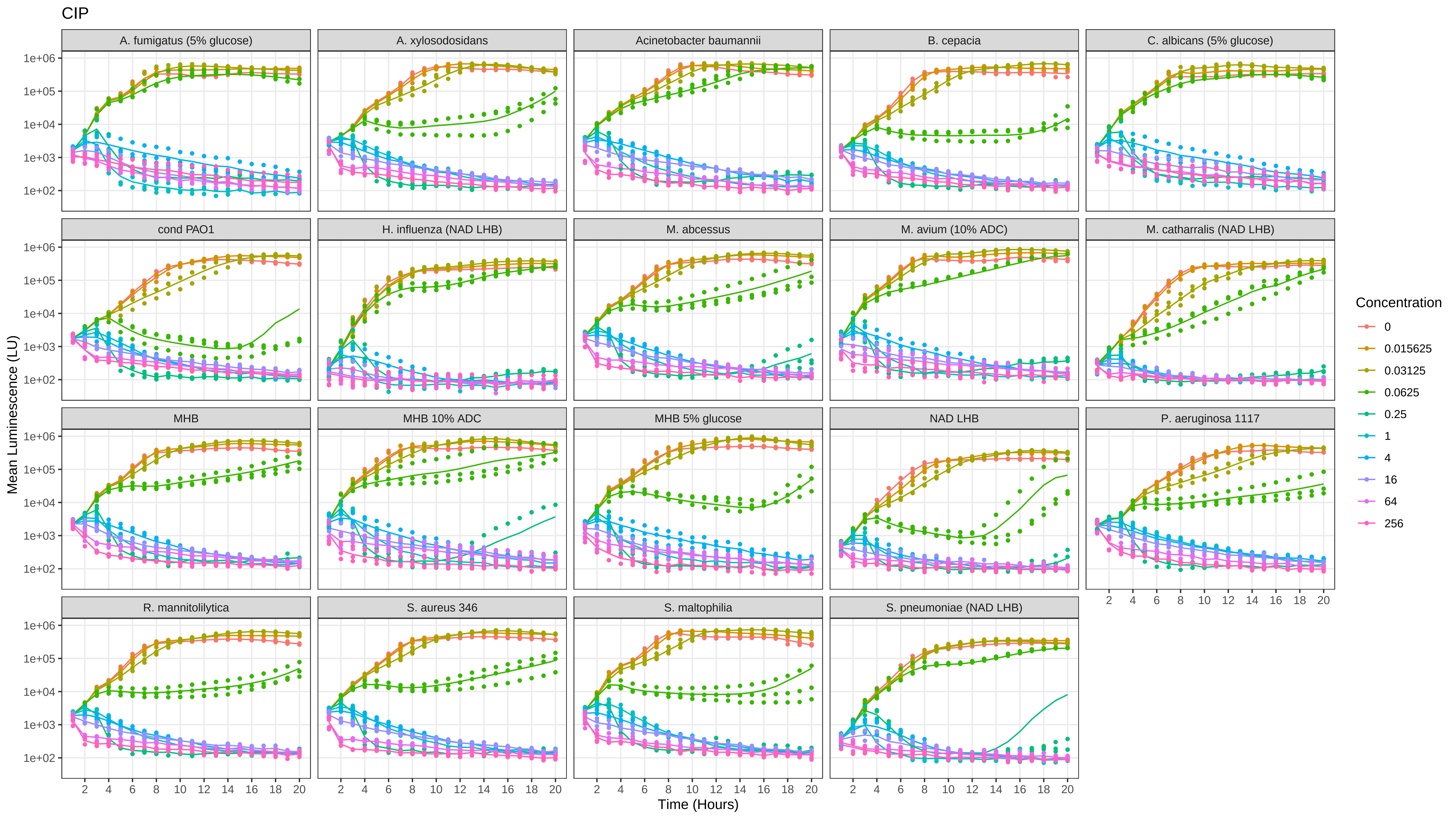

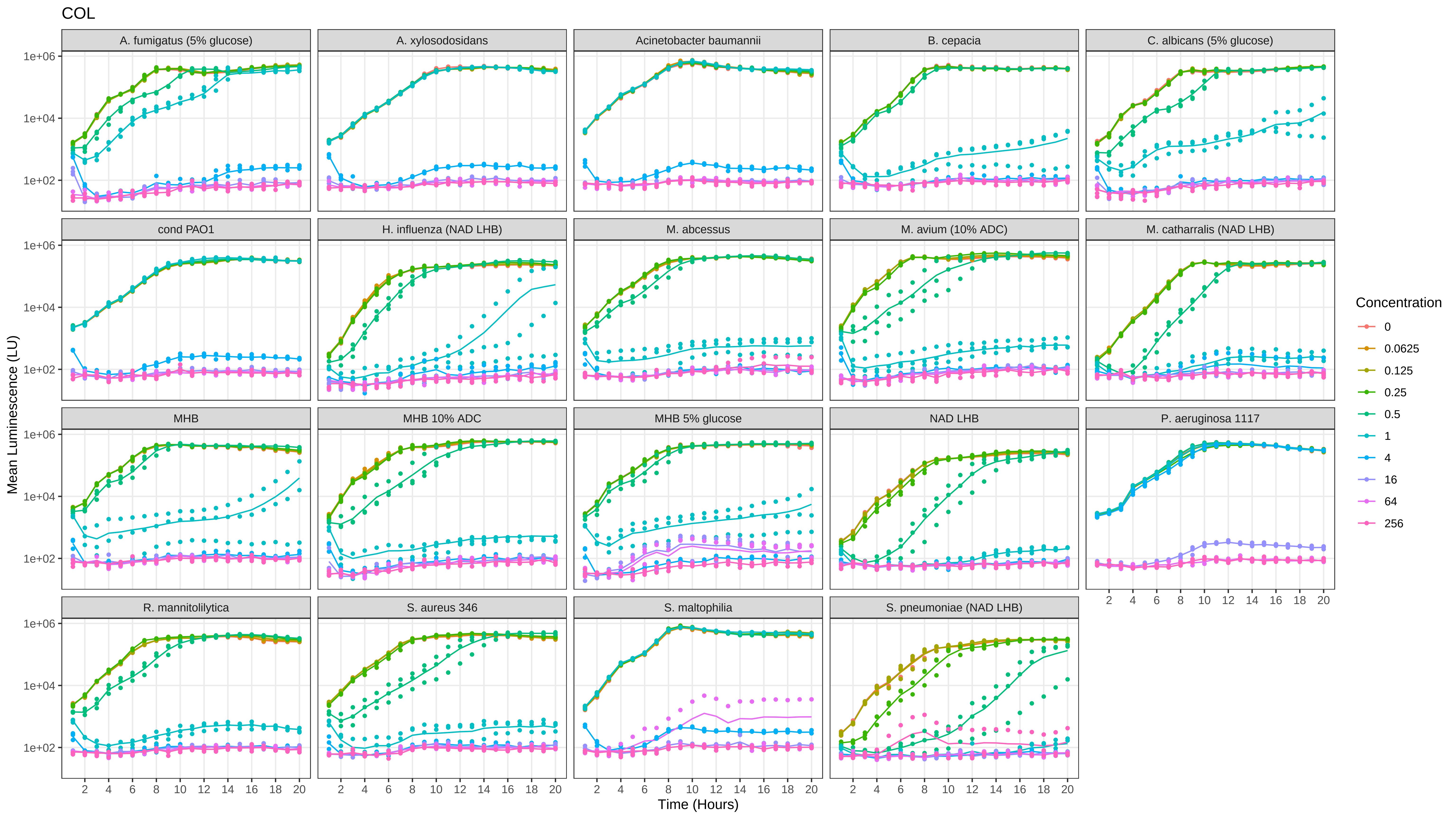

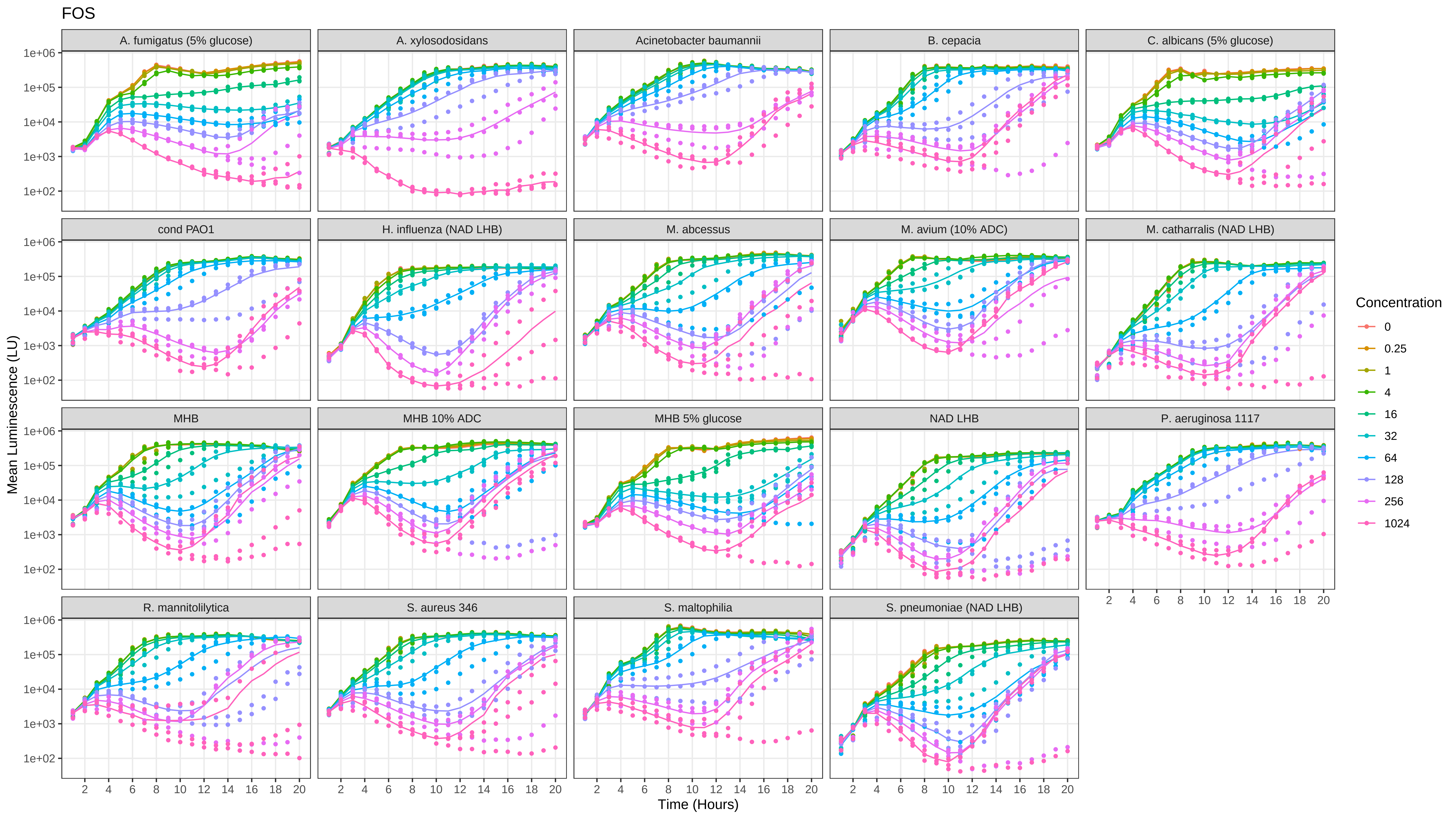

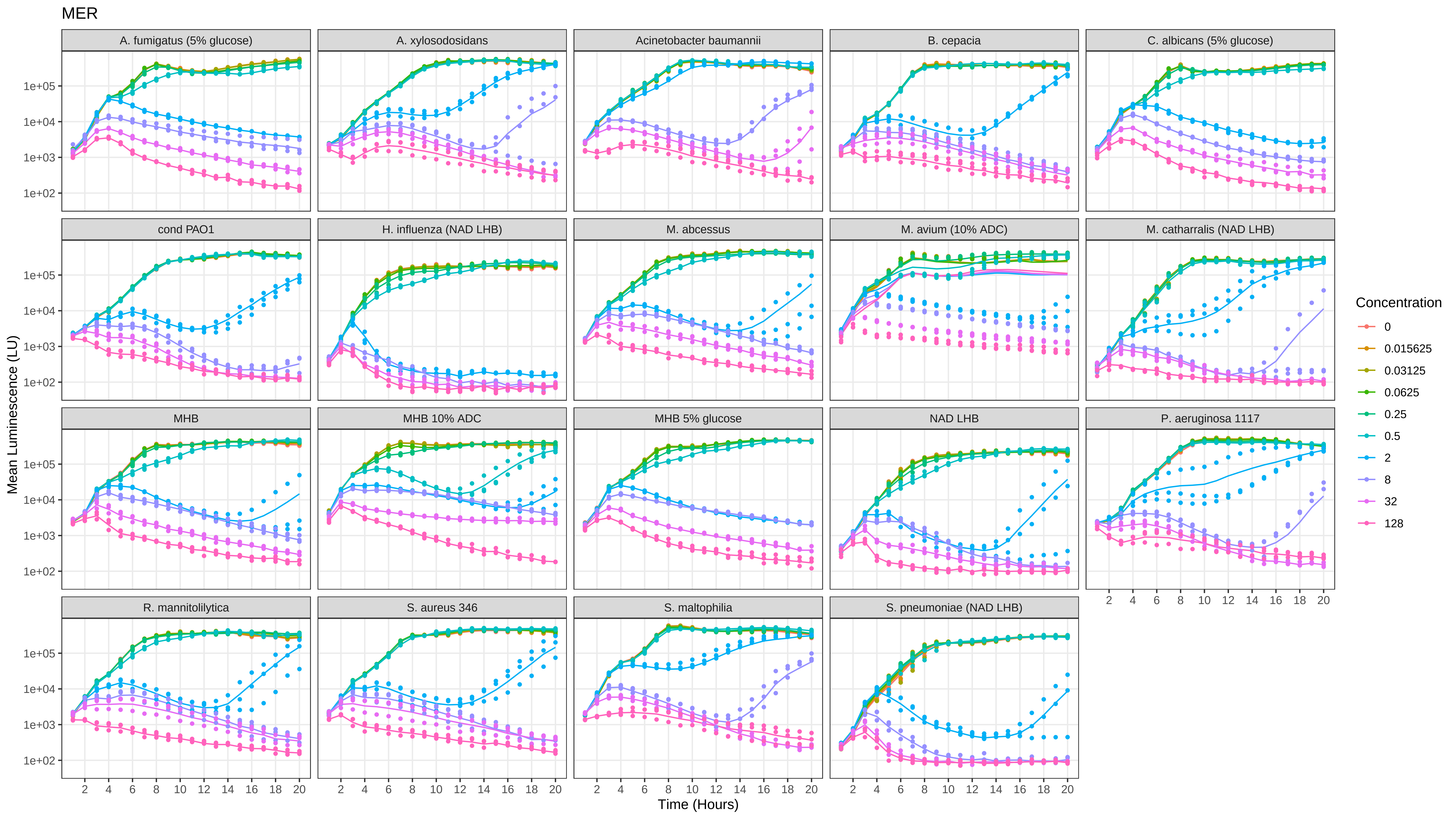

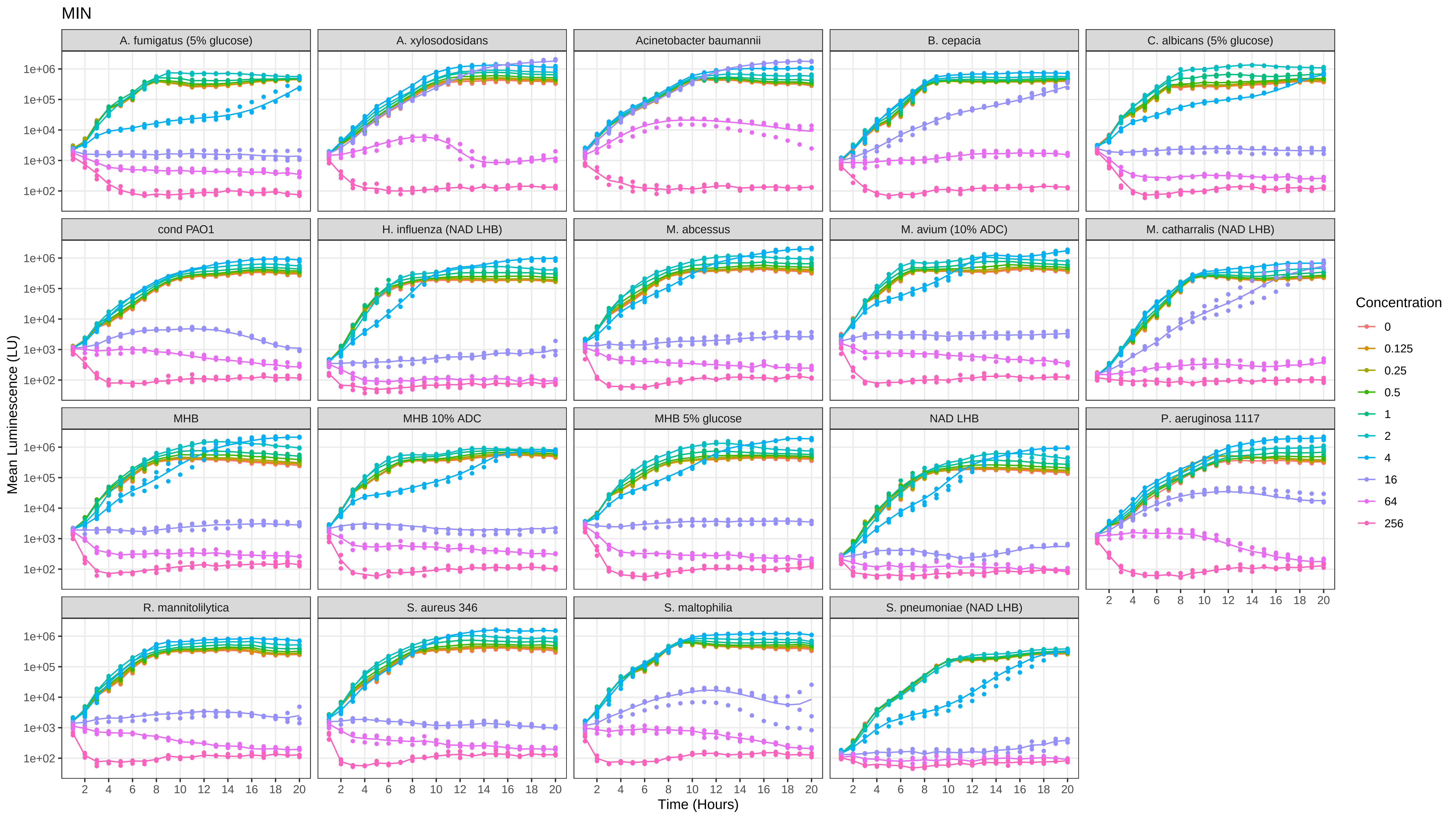

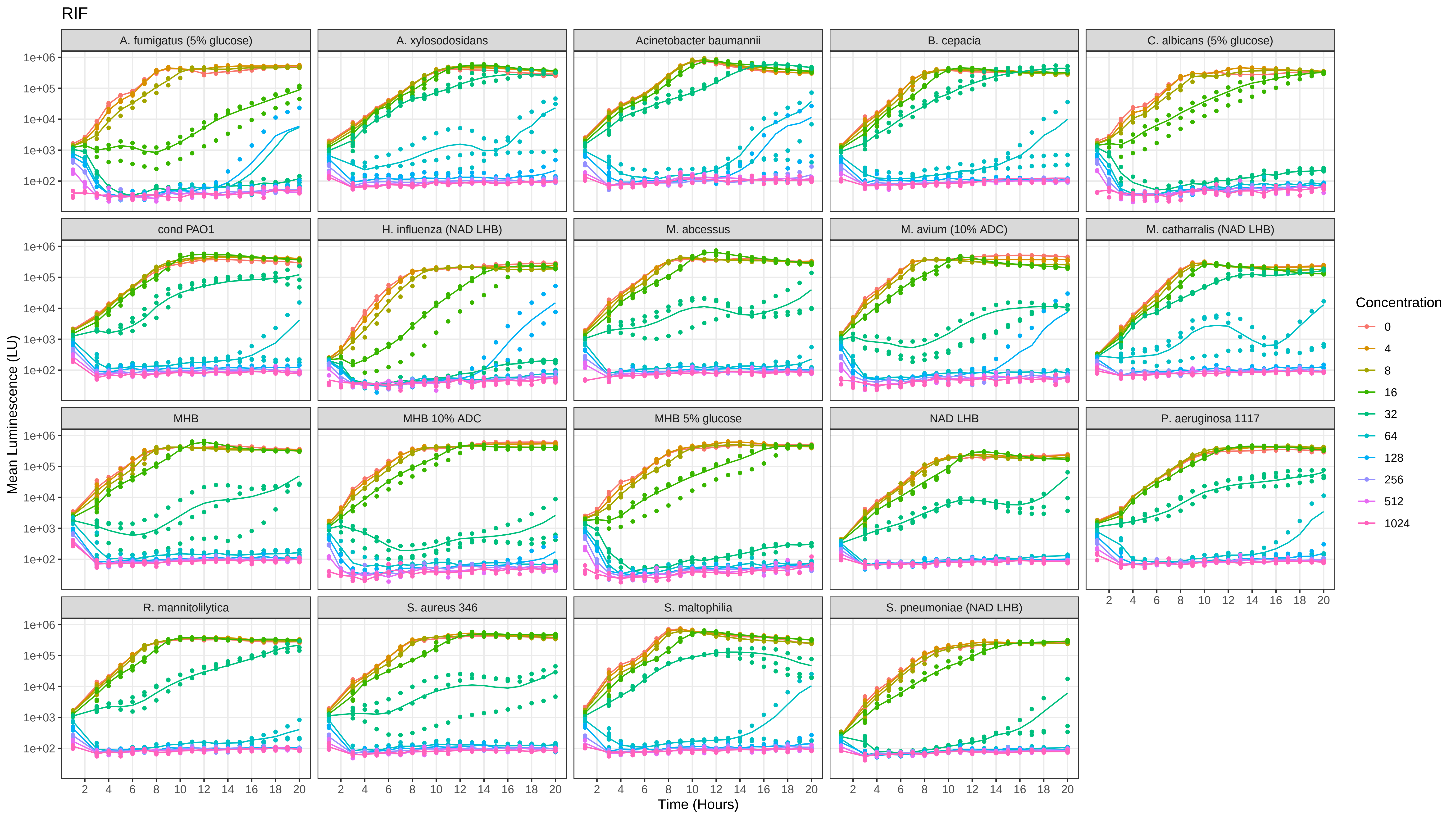

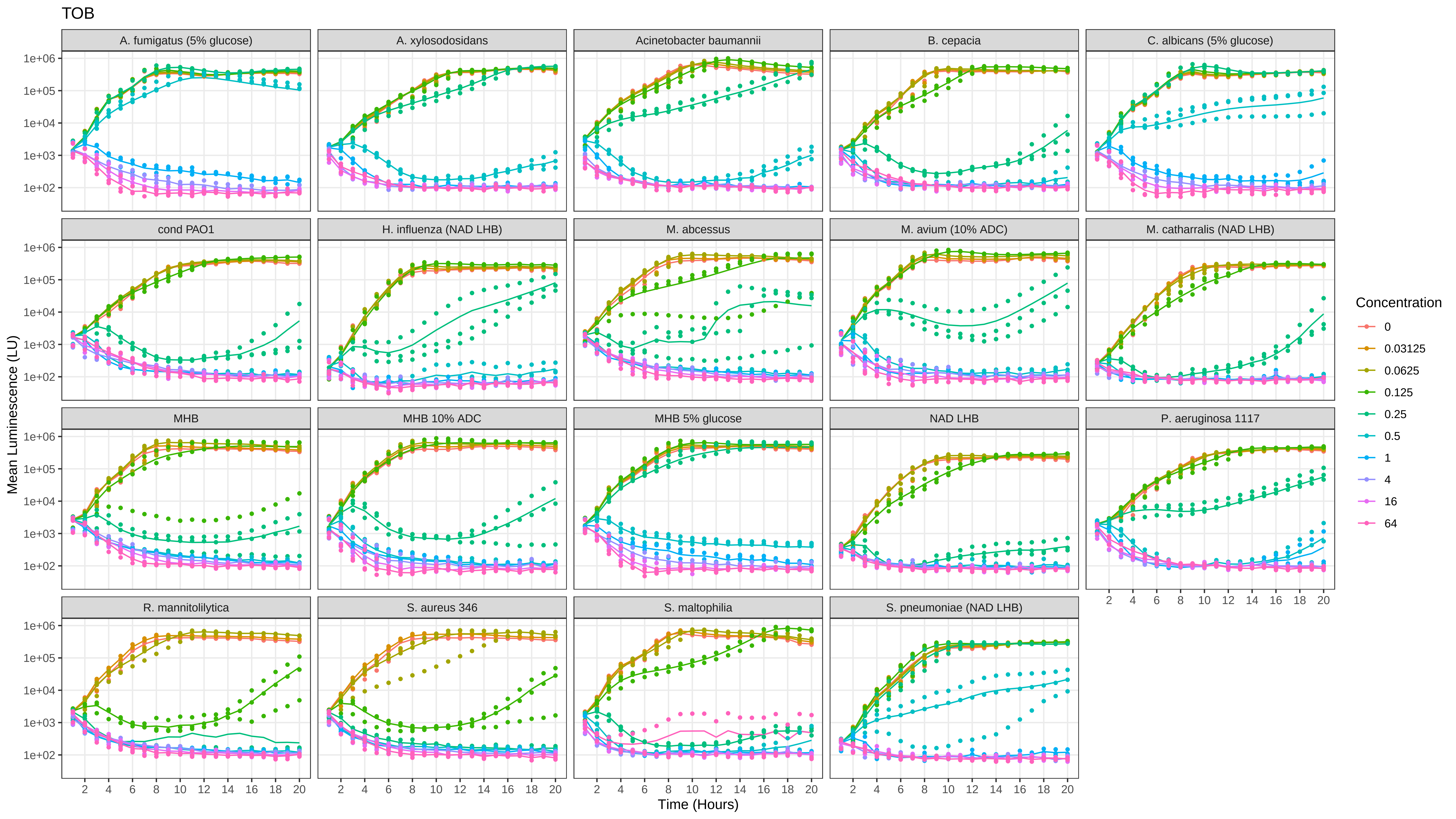

### SI_8_concentration_effect_curves.pdf

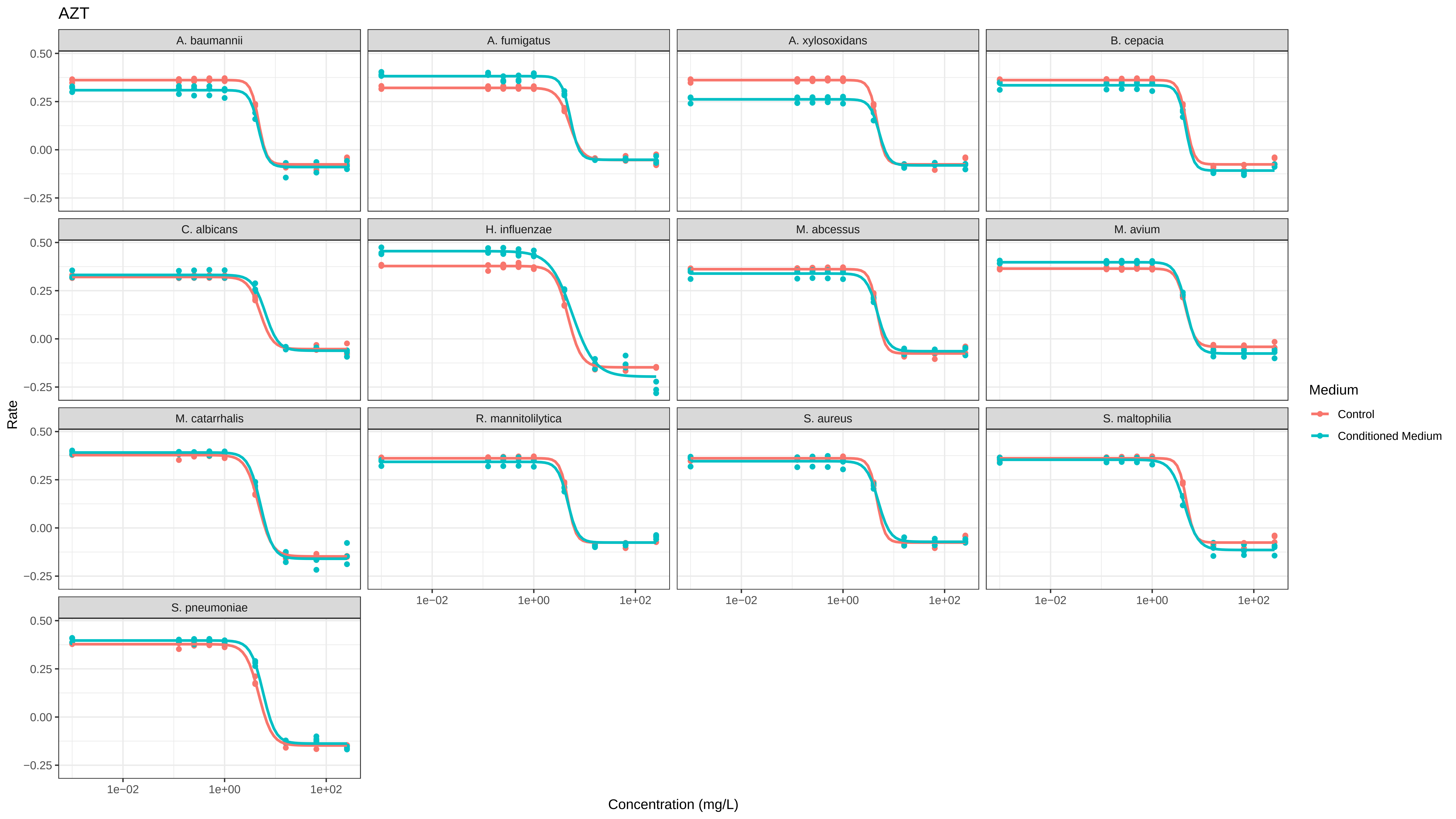

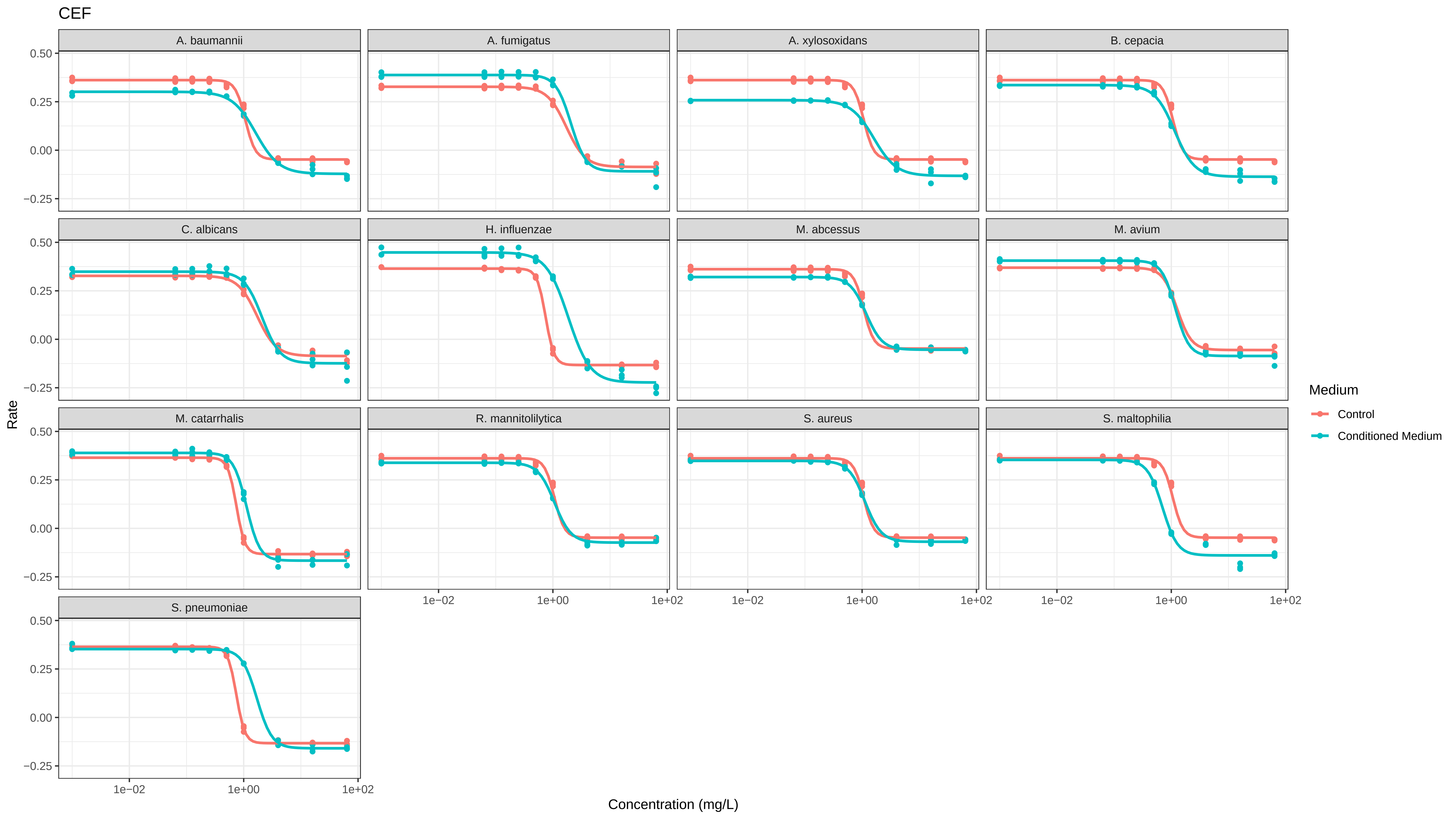

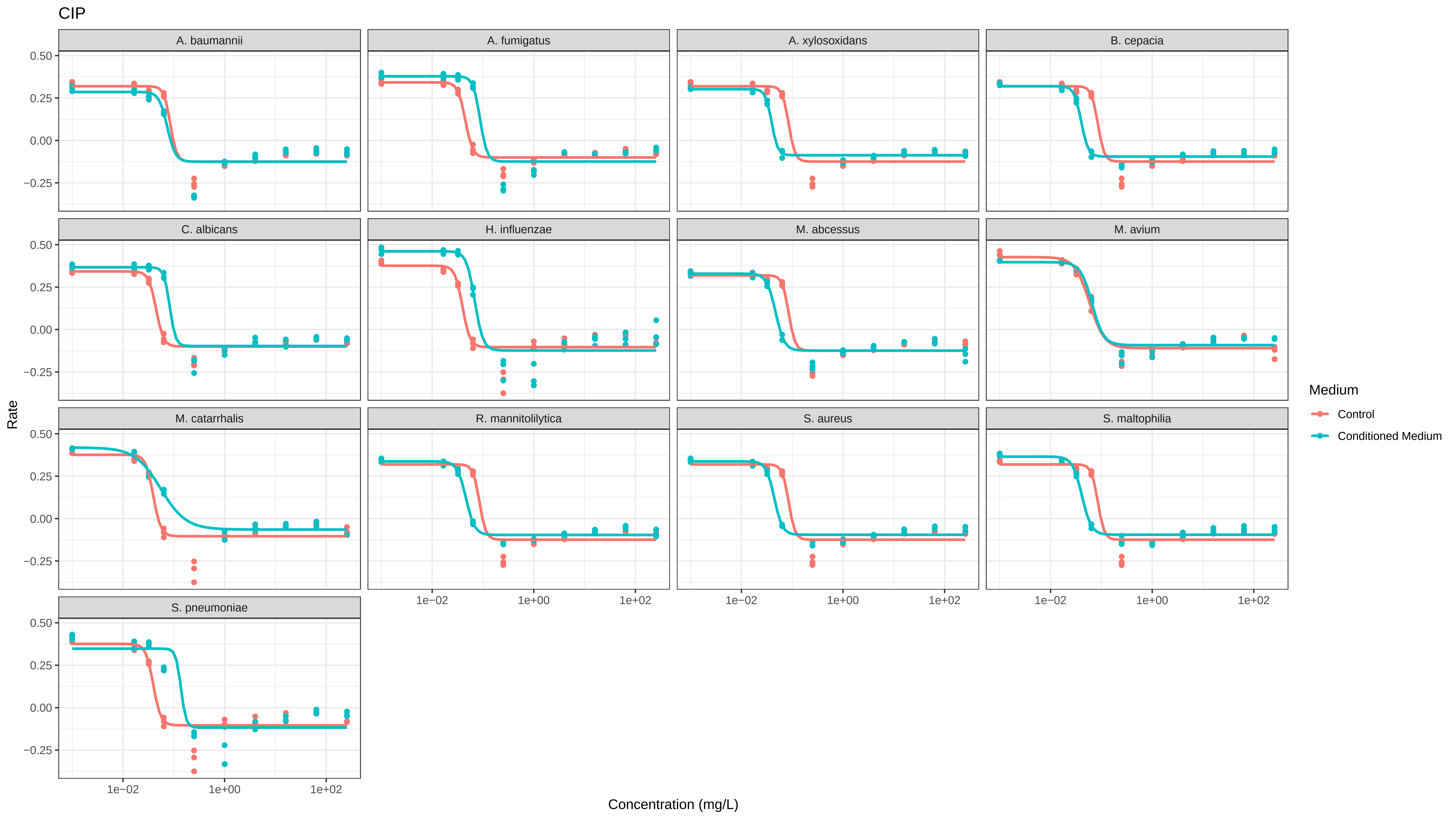

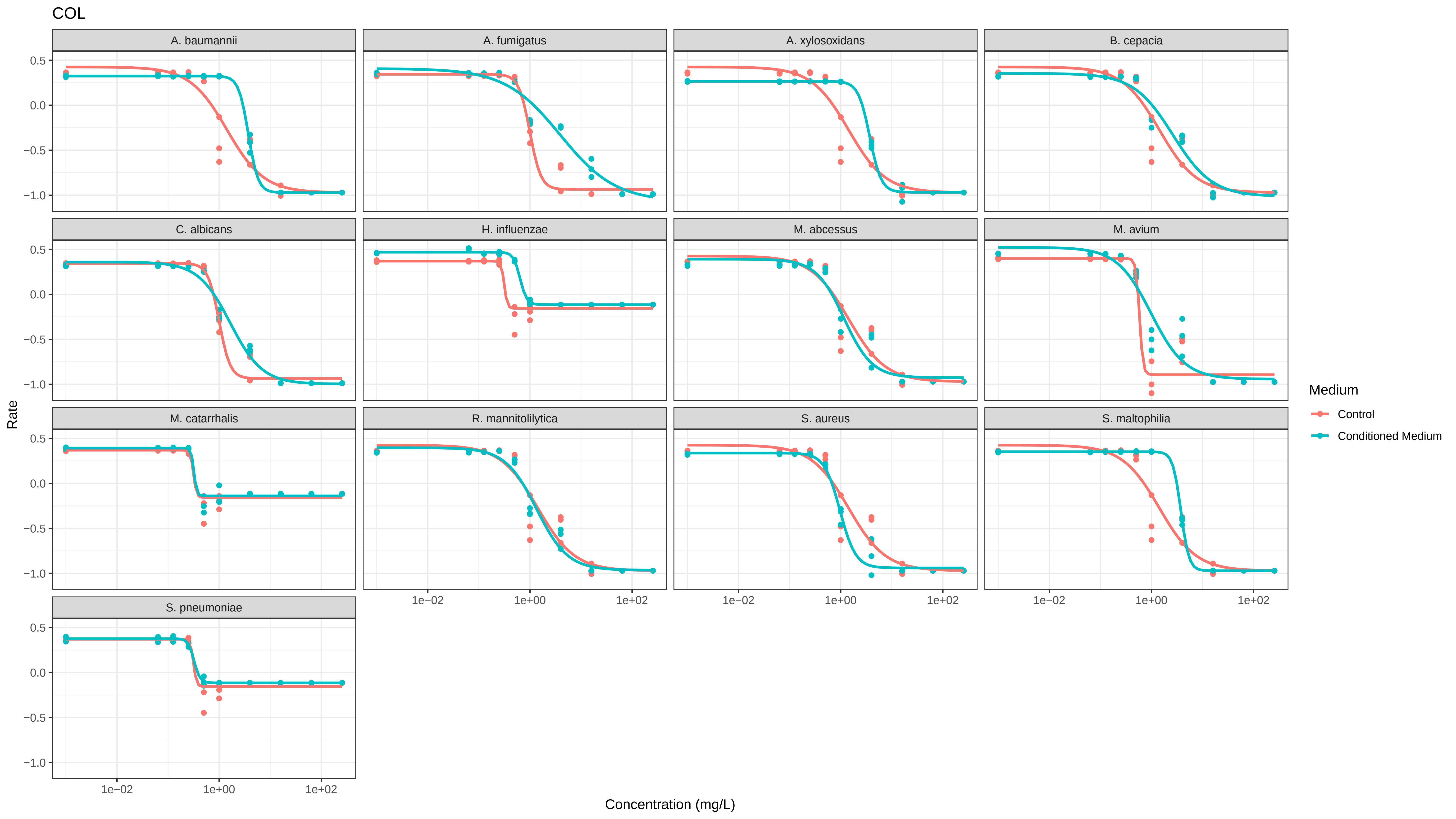

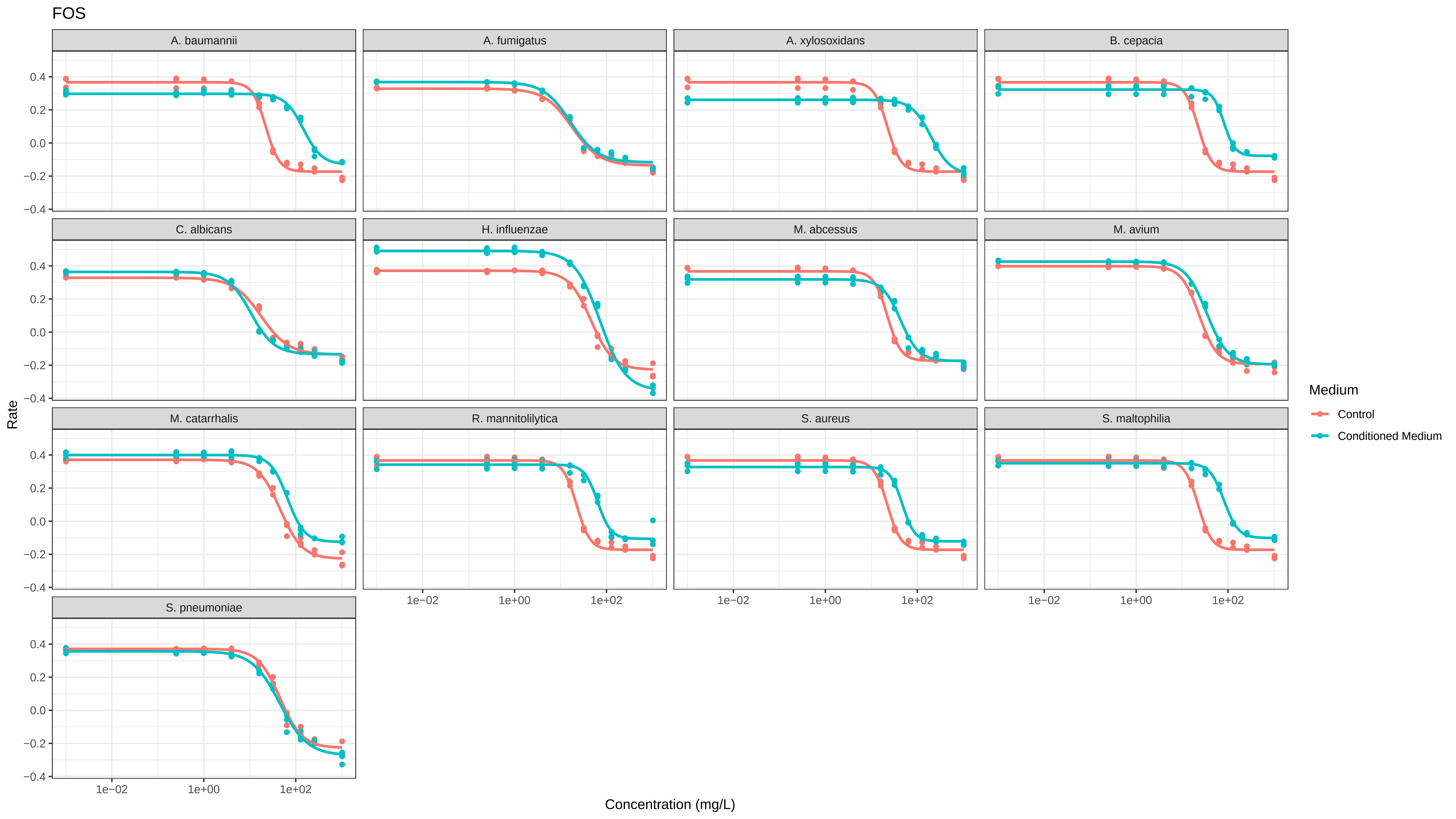

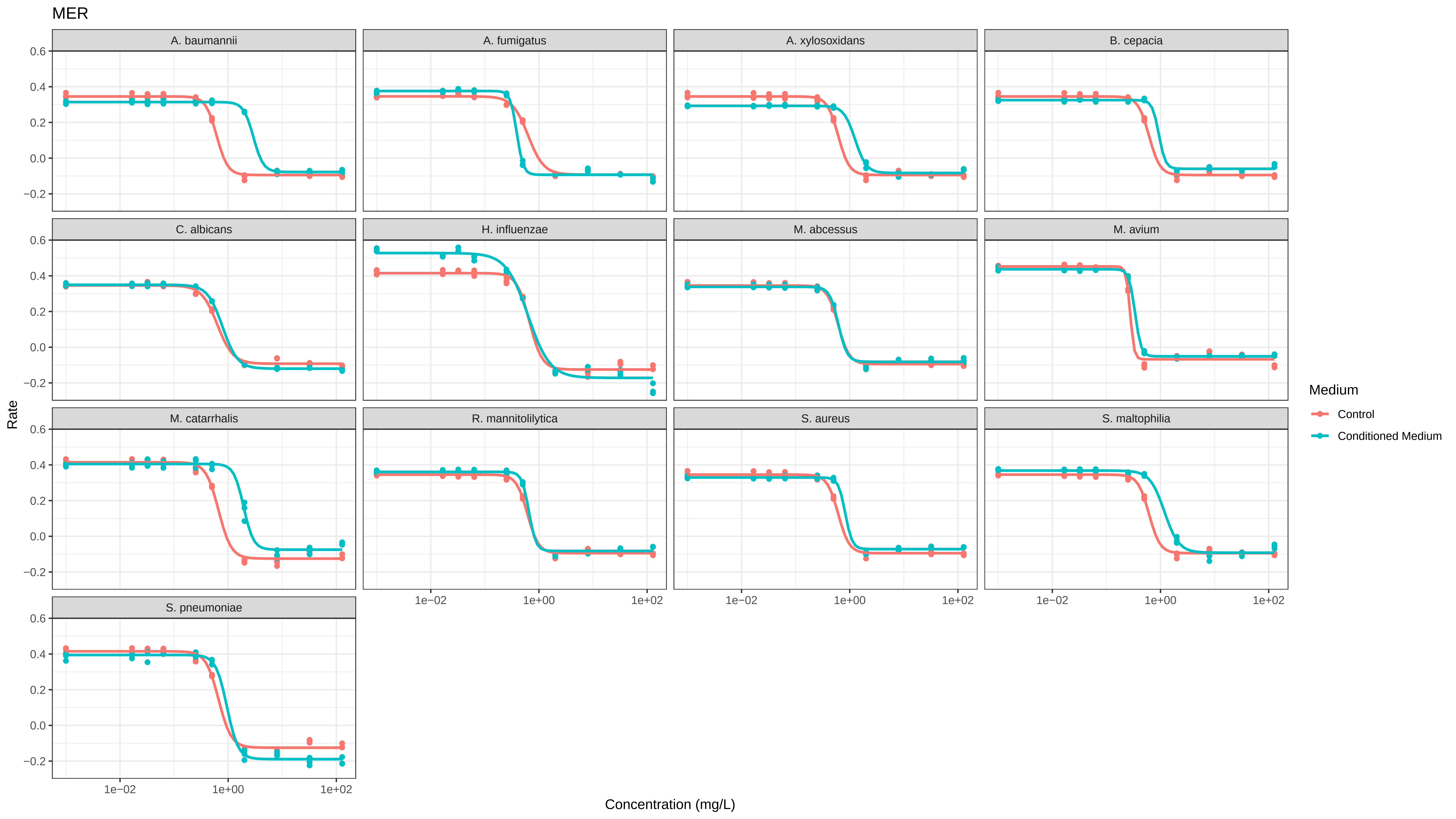

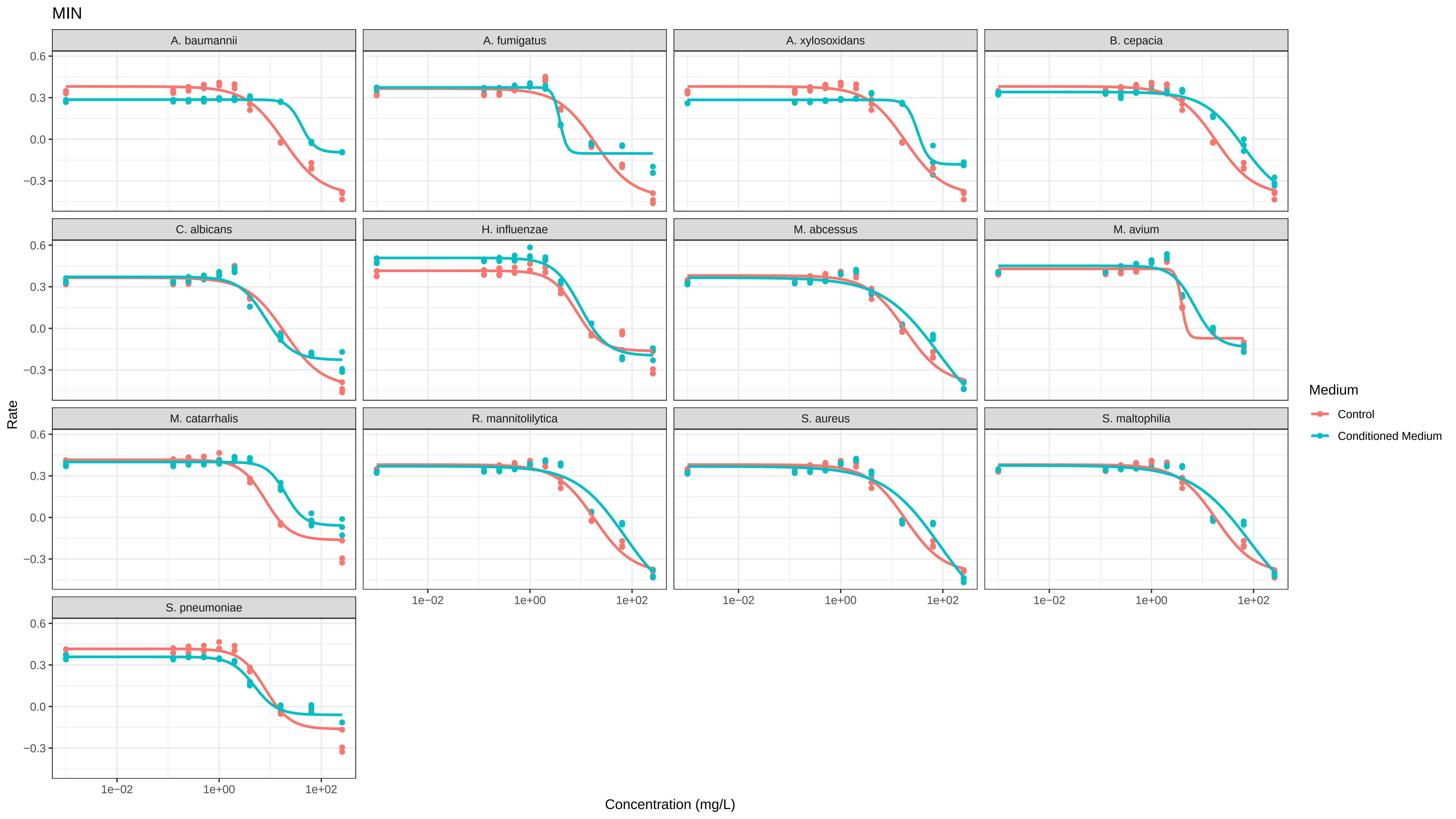

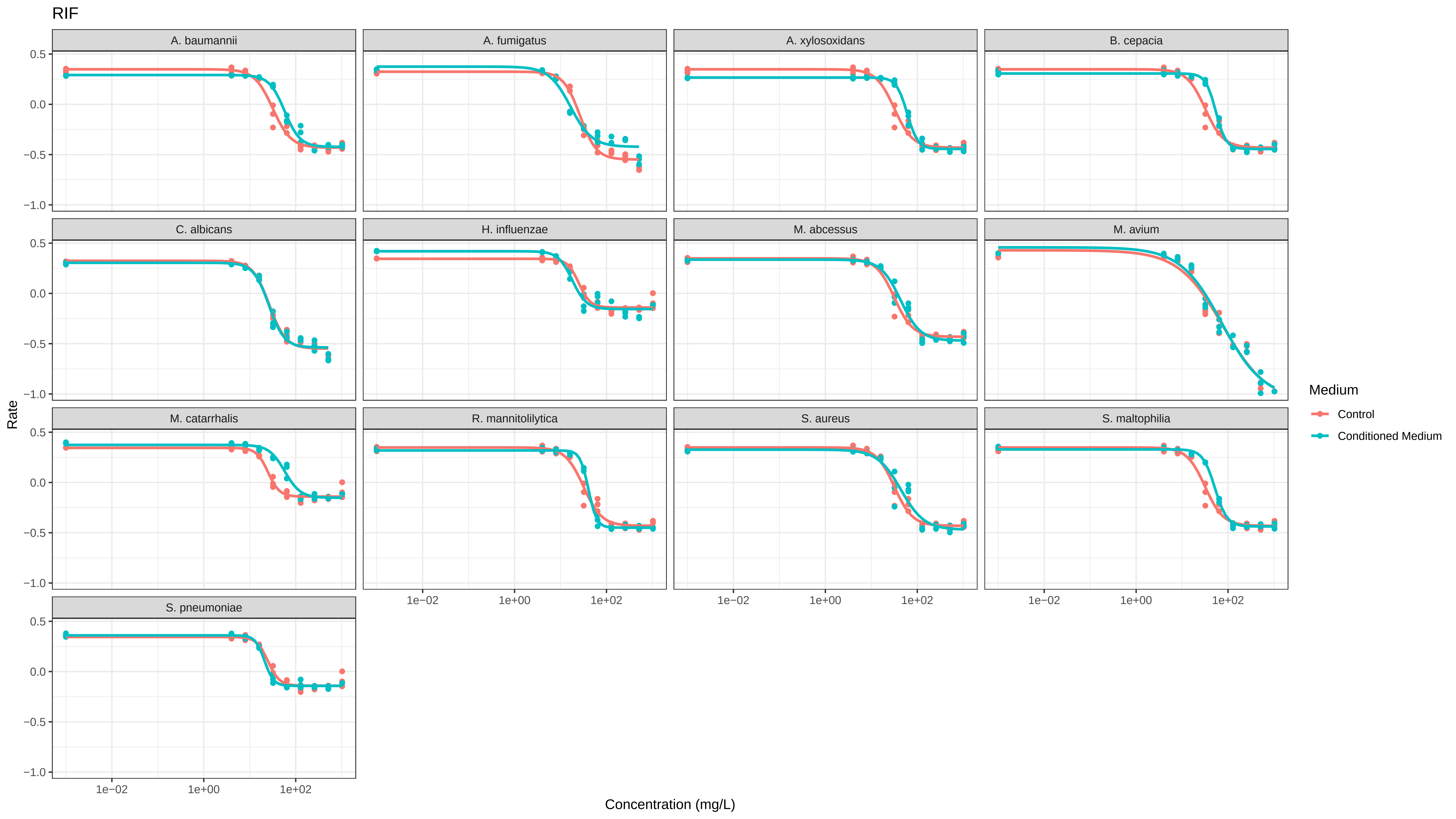

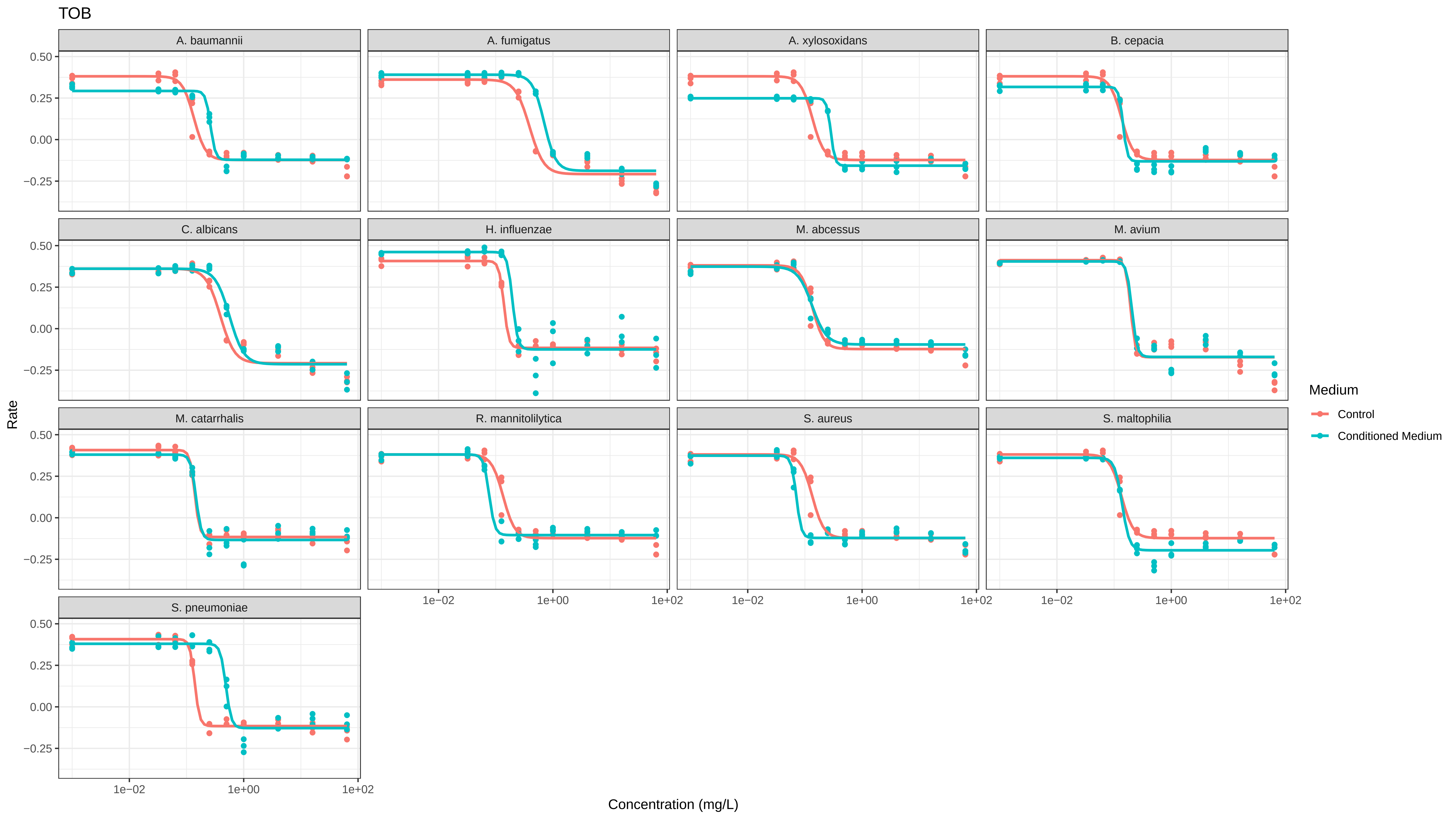

### SI_9_Distribution_PD_Parameters.pdf

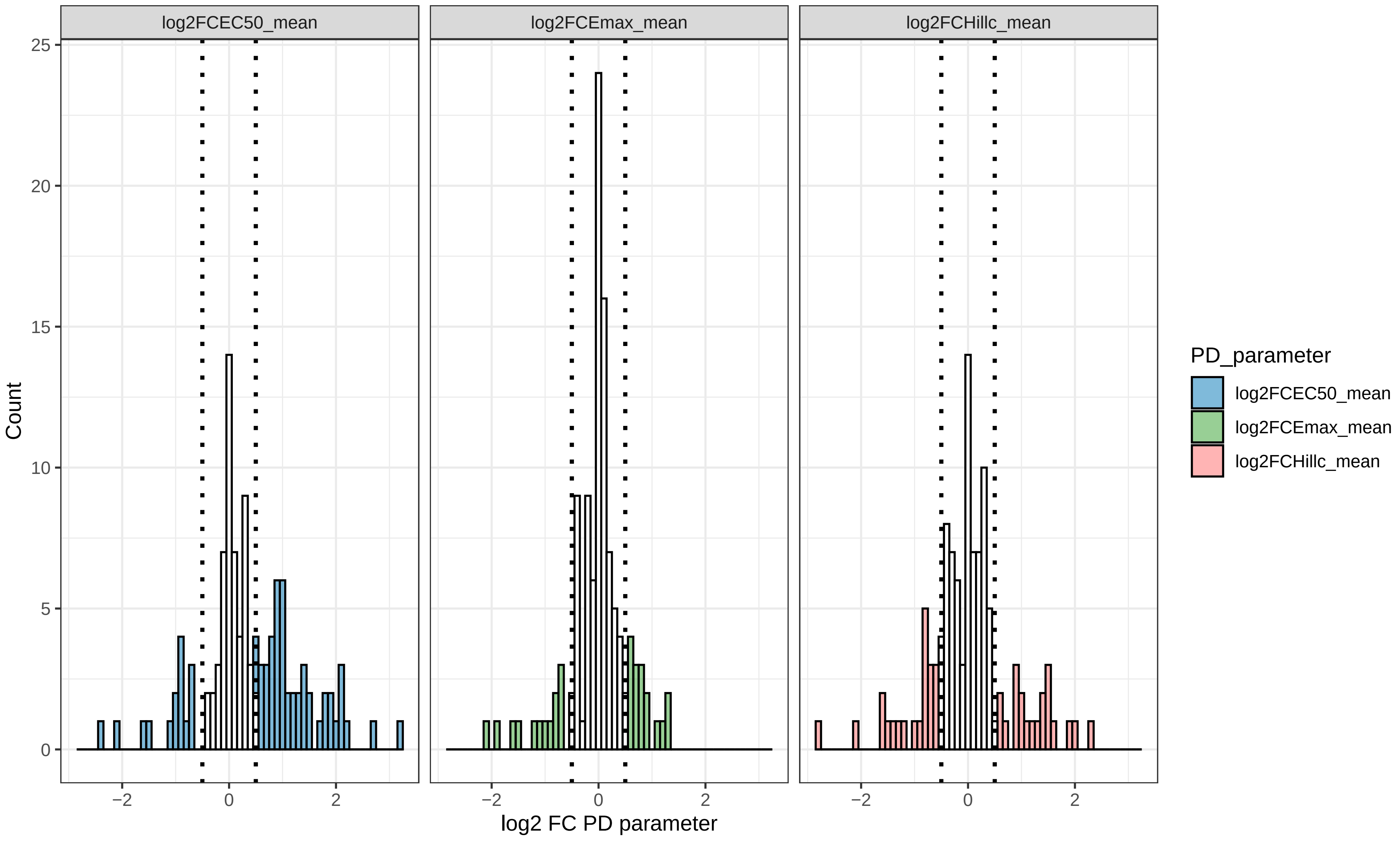
